# Stress-associated purinergic receptors code for fatal suicidality in the hippocampal-hypothalamic-prefrontal circuit

**DOI:** 10.1101/2022.11.22.516142

**Authors:** Lin Zhang, Ronald W.H. Verwer, Joop van Heerikhuize, Rawien Balesar, Felipe Correa-da-Silva, Zala Slabe, Paul J. Lucassen, Dick F. Swaab

**Affiliations:** Neuropsychiatric Disorders Lab, Neuroimmunology Group, Netherlands Institute for Neuroscience, an Institute of the Royal Netherlands Academy of Arts and Sciences, Amsterdam, the Netherlands; Department of Endocrinology and Metabolism, Amsterdam Gastroenterology Endocrinology and Metabolism, Amsterdam University Medical Center (UMC), University of Amsterdam, Amsterdam, the Netherlands; Laboratory of Endocrinology, Amsterdam University Medical Center (UMC), University of Amsterdam, Amsterdam, the Netherlands; Faculty of Medicine, Institute of Pharmacology and Experimental Toxicology, University of Ljubljana, Ljubljana, Slovenia; Brain Plasticity Group, Faculty of Science, Swammerdam Institute for Life Sciences, University of Amsterdam, Amsterdam, the Netherlands

**Keywords:** Suicide, legal euthanasia, hippocampus, hypothalamus, prefrontal cortex, purinergic receptors

## Abstract

Imbalanced purine metabolism is a key neurological basis for suicide and mood disorders (MD), wherein purinergic receptors in stress-sensitive cerebral regions are thought to be differentially activated. A hippocampal network that links the hypothalamus and prefrontal cortex implements an affective sensation of stress. We discovered that the hippocampus encoded fatal suicidal ideations in the dentate gyrus (DG) by a considerable amount of the granule cell nuclei with P2X purinoceptor 7 (P2RX7) expression, irrespective of the underlying MD. Compared to controls, patients with MD showed microglial dyshomeostasis throughout the hippocampal formation. Strikingly, P2Y purinoceptor 12 (P2RY12)-expressing microglia with segmented processes were remarkably present in the superficial layers of the medial entorhinal cortex (mEnt) in individuals with fatal suicidality. In the hypothalamic stress-sensitive nuclei, P2RY12^+^ microglia were more expressed in the supraoptic nucleus in MD and even higher when fatal suicidality was present. In the prefrontal cortex, P2RX7 transcripts sharply dropped in suicidal individuals, possibly removing the prefrontal inhibition of the hippocampus and hypothalamus. Confounder analysis showed that the suicide-specific molecular features faded when the postmortem delay was prolonged. Our findings imply that fatal suicidality presents with unique neuropathological alterations. The DG and mEnt are two crucial areas for deciphering the suicidal consequences. By including brain samples from legal euthanasia donors, suicide-specific biosignatures can be maximally retained. Decoding the bioactive framework through key genes, brain regions and neurological processes involved in suicide neuropathology may provide novel therapeutic strategies for suicidal individuals who are beyond the reach of mental health care.

## Introduction

Suicide is a leading cause of death worldwide and has significant emotional, physical, and economic impact on sufferers and their relatives ^1^. It frequently occurs in the context of a mood disorder (MD), but these two conditions have at least partially distinct molecular signatures ^2^. Disturbed purine metabolism is a key component in the pathophysiology of suicide ^3–7^. A large-scale pharmacoepidemiologic study has shown that the prescription of folic acid, which plays a key role in purine biosynthesis, is linked to lower rates of suicide attempts in adults ^8,9^. Measuring the expression of purinergic receptors can predict both extracellular purine activities and intracellular biological responses in suicide and MD. One potent candidate is P2X purinoceptor 7 (P2RX7), an adenosine triphosphate (ATP)-gated ion channel, which is implicated as a novel therapeutic target for depression ^10^. Human genetic studies have linked the highly polymorphic *P2RX7* gene to MD ^11^. In peripheral blood mononuclear cells of suicide completers, P2RX7 mRNA levels were lower as compared to non-suicidal controls ^12^. The other is a microglia-expressed adenosine diphosphate (ADP) chemoreceptor, P2Y purinoceptor 12 (P2RY12), which is found to be more expressed in the prefrontal cortex (PFC) of suicide completers ^4–7^. In addition to the PFC, animal models of depressive symptoms have further shown disturbed purine metabolism in the hypothalamus and hippocampus ^13,14^.

The hippocampus, which is incorporated in a limbic-cortical-hypothalamic circuit, has been functionally implicated in the connection between stress and cognition. Whole-brain functional neuroimaging studies have shown that hippocampal connectivity with the hypothalamus and PFC specifically predicts the subjective sensation of stress in humans ^15^. While studies have discrepancies in summarizing brain circuits associated with suicidal behaviors ^16–18^, there are no follow-up studies that focus on people who later died by suicide, which is essential for studying the neurocircuitry of strong and determined suicide intent. One possibility is that intense and predetermined suicide tendencies (fatal suicidality) would be better detectable than transient or sporadic risks (non-fatal suicidality) by imaging-based indicators. Therefore, biological insights into suicide are more likely to be implanted in the candidate circuits of people with fatal suicidality.

To address the issue, we collected 321 samples (including 77 hippocampi, 44 hypothalami, and 200 PFCs) from donors who had MD, suicidality or accomplished suicide, and their matched controls. We examined the transcriptional and translational alterations of stress-related purinergic receptors (P2RX7 and P2RY12), as well as two glial markers (GFAP and CD68) to probe the neuroinflammatory and microglial phagocytic implications in suicide and MD. In contrast to the earlier measurements that were carried out in bulk tissue preparation, which might have overlooked the specific role of selective subregions, we quantified five subdivisions in the hippocampal formation separately (dentate gyrus (DG), *cornu ammonis* (CA) 1, CA2-4, subiculum (Sub) and entorhinal cortex (Ent)), and two stress-sensitive hypothalamic nuclei (supraoptic nucleus, SON and paraventricular nucleus, PVN). Based on our previous studies ^5,6^, we examined P2RX7 transcripts in the dorsolateral prefrontal cortex (DLPFC) and anterior cingulate cortex (ACC), to address the question of whether P2RX7 expression profiles were regionally modulated by completed suicide or coexistent psychiatric diagnoses.

## Materials and methods

### Human brain materials

#### Hippocampus

Seventy-seven paraffin-embedded tissue blocks containing the anterior or middle hippocampus were obtained from the Netherlands Brain Bank (NBB, Amsterdam, the Netherlands, Director: Prof. Dr. Inge Huitinga). The donors or their next of kin provided permission for the autopsy, use of brain material, and clinical documentation for research. The samples were divided into two main groups: 56 MDs and 21 matched non-psychiatric controls (Ctr) without a neurological disorder. There were three subsets in the MD group and two subsets in the Ctr group. The MD group contained 10 subjects with MD who completed suicide (MDS), 21 subjects with MD and fatal suicidality that led to legal euthanasia (MDE), and 25 subjects with MD who died of natural causes (MDN). For this cohort, we carefully selected a matched group that contained 10 subjects who displayed fatal suicidality and died of legal euthanasia for incurable diseases (CE), and 11 subjects who died of natural causes (CN). According to the suicidal consequences, subjects were regrouped into three subsets. Twenty-three subjects who did not report suicidality lifelong and died of natural causes (NS, no suicidality), 13 subjects who had a clinical history of suicide tendency but died of natural causes (NFS, non-fatal suicidality), and 41 subjects who died of completed suicide or legal euthanasia (FS, fatal suicidality). After we introduced the classifications of suicidality into grouping, which was documented as suicidal ideations and suicide attempts, subjects were subdivided into non-fatal suicidal ideation (NFSI) and suicide attempts (NFSA), and fatal suicidal ideations (FSI) and suicide attempts (FSA).

#### Hypothalamus

Forty-four tissue blocks containing the entire hypothalamus were obtained from the NBB according to the same procedure as described above. Subjects were collected and divided into two main groups: 28 MDs and 16 Ctrs. The MD group contained 10 MDS, 7 MDE, and 11 MDN subjects. The Ctr group contained 5 CE and 11 CN subjects.

#### Prefrontal cortex

Two hundred RNA samples, together with their demographic information and medical data, were obtained from the Depression and Array Collections of the Stanley Medical Research Institute (SMRI, Bethesda, MD, the USA, Director: Dr. Maree J. Webster). The SMRI provided us with RNA from the isolated grey matter of the DLPFC (Brodmann area 46) and ACC (Brodmann area 24). They were obtained from patients with MD (N = 54; major depressive disorder, MDD N = 24; bipolar disorder, BD N = 30) and matched controls (Ctr, N = 36) without a history of suicidal behaviors or any major psychiatric diagnosis. The cause of death for 30 of 54 patients with MD was suicide. Twenty-nine patients with MD had a history of psychotic features. The other cases and all control subjects died of natural causes or accidents. We distinguished the following subsets within the MD group: 1) patients with MDD or BD; 2) patients with MDD/BD who died of suicide (MDDS/BDS) or non-suicidal causes (MDDNS/BDNS); and 3) patients with or without a history of psychotic features (MDDP/BDP or MDDNP/BDNP).

Clinical details of every donor can be found in the <u>Supplementary Information (**Table S1**)</u>.

### Characterizing and counting P2RX7^+^ nuclei in the hippocampal formation

First, the density of the granule cells in the DG of every section was determined to assess structural stability. Second, P2RX7 and GFAP immunohistochemical staining was performed to investigate their expression in the hippocampal subfields. The distribution of P2RX7^+^ nuclei and P2RX7^+^ astrocytes, and the total expression of P2RX7 and GFAP in the hippocampus were determined and their associations with suicide and MD were established. Bright-field images were acquired by a ZEISS Axio Scanner and analyzed using QuPath (see <u>Supplementary Information</u>).

### Recalling microglial functions and morphological profiles in the hippocampus

P2RY12 and CD68 immunohistochemical and immunofluorescent staining were performed as markers for microglial homeostatic and phagocytic capacity. Bright-field images were analyzed using QuPath and ImageJ for expression and morphological measurements. Fluorescent images were captured by a Leica laser-scanning microscope SP8 and processed with Imaris for 3D reconstruction. The density and radius of the microglial cell body, the size of the microglial cell body with processes and fragments (microglial segments that were disconnected from the cell body) were determined (see <u>Supplementary Information</u>).

### Profiling the expression of purinergic receptors and glia in the hypothalamic nuclei

First, the volumes of the hypothalamic SON and PVN were determined based on the thionine staining. Second, P2RY12, CD68, P2RX7 and GFAP immunohistochemical staining were performed to investigate their expression in the SON and PVN. Images were analyzed with Image-Pro. CD68 and P2RY12 immunofluorescent staining and image processing were performed with the same protocols mentioned above (see <u>Supplementary Information</u>).

### Transcriptional analysis of P2RX7 expression in the PFC

cDNA synthesis of the PFC-RNA was performed for real-time qPCR studies (see <u>Supplementary Information</u>). The transcriptional data of prefrontal P2RY12, GFAP, and CD68 expression in relation to suicide and MD have been included in our previous publications ^5,6^.

### Statistical analysis

S+ software (version 8.2, TIBCO, Seattle, WA, USA) was used for statistical analysis.

The Chi-square test was used for the analysis of categorical data (suicidal classification and gender). For interval data, the Mann-Whitney test (2 samples) or the Kruskal-Wallis test with multiple comparisons (more than 2 samples) was used ^19^. Before processing real-time qPCR data, the values were ^10^log-transformed to enable simple reference gene correction and conventional statistical procedures. The reason for this transformation was that the observed Ct values used in order to quantify gene expression are exponents of the PCR efficiency. Application of the log-transformation yields an additive statistical model and, after all, statistical procedures have been finished, the data are back-transformed and presented as fold-changes. In the case of multiple testing, a Benjamini-Hochberg correction ^20^ of *p*-values was applied. When the Kruskal-Wallis test was used in combination with the Benjamini-Hochberg correction, we proceeded in a 2-step way. As multiple comparisons in the Kruskal-Wallis test are only allowed if the global *p* < 0.05 ^19^, we first corrected the global *p*-values and then selected for further analysis only those biomarkers for which this requirement was met. For each appropriate comparison, the corresponding *p*-values were pooled and corrected according to Benjamini-Hochberg. All tests were 2-sided. Figures were prepared with GraphPad Prism (statistical analysis) or BioRender (illustrative graphs).

## Results

### A prevalence of P2RX7-positive nuclei in the hippocampal formation mark fatal suicidality

In the hippocampus of individuals who died of natural causes, P2RX7 was generally expressed by glial cells (**Fig. 1d** and **1l**, **S1d** and **S1h**, and **S2a** and **S2i**). In the DG, granule cell densities were similar between these groups (**Fig. 1c**). Strikingly, in individuals who died by suicide or legal euthanasia, a considerable number of granule cells expressed P2RX7 within their nuclei (**Fig. 1e-g**). Characteristic features of the P2RX7^+^ nuclei were their chromatin condensation and marginalization (**Fig. 1e-g**). To qualify this finding, we determined the percentage of over 5% of P2RX7^+^ nuclei within the total granule cells to be a proportioned positive subject, in which P2RX7^+^ nuclei were at detectable levels in microscopy practice. The donors who died by legal euthanasia or suicide had much higher proportions than those who died of natural causes (**Fig. 1h**). Quantitative results presented similar enhancements in the ratio of P2RX7^+^ nuclei to the total number of nuclei in the granule cell layer (CE vs CN: *p* = 0.0008, MDE vs MDN: *p* < 0.0001, MDS vs MDN: *p* = 0.0008, MDS vs CN: *p* = 0.014, MDE vs CN: *p* = 0.0002; **Fig. 1i**). No differences were found between MD and controls following legal euthanasia (MDE vs CE: *p* = 0.74), or in natural death cases (MDN vs CN: *p* = 0.35). In MD, suicide cases did not show alterations compared to legal euthanasia (MDS vs MDE: *p* = 0.71).

**Figure 1.**
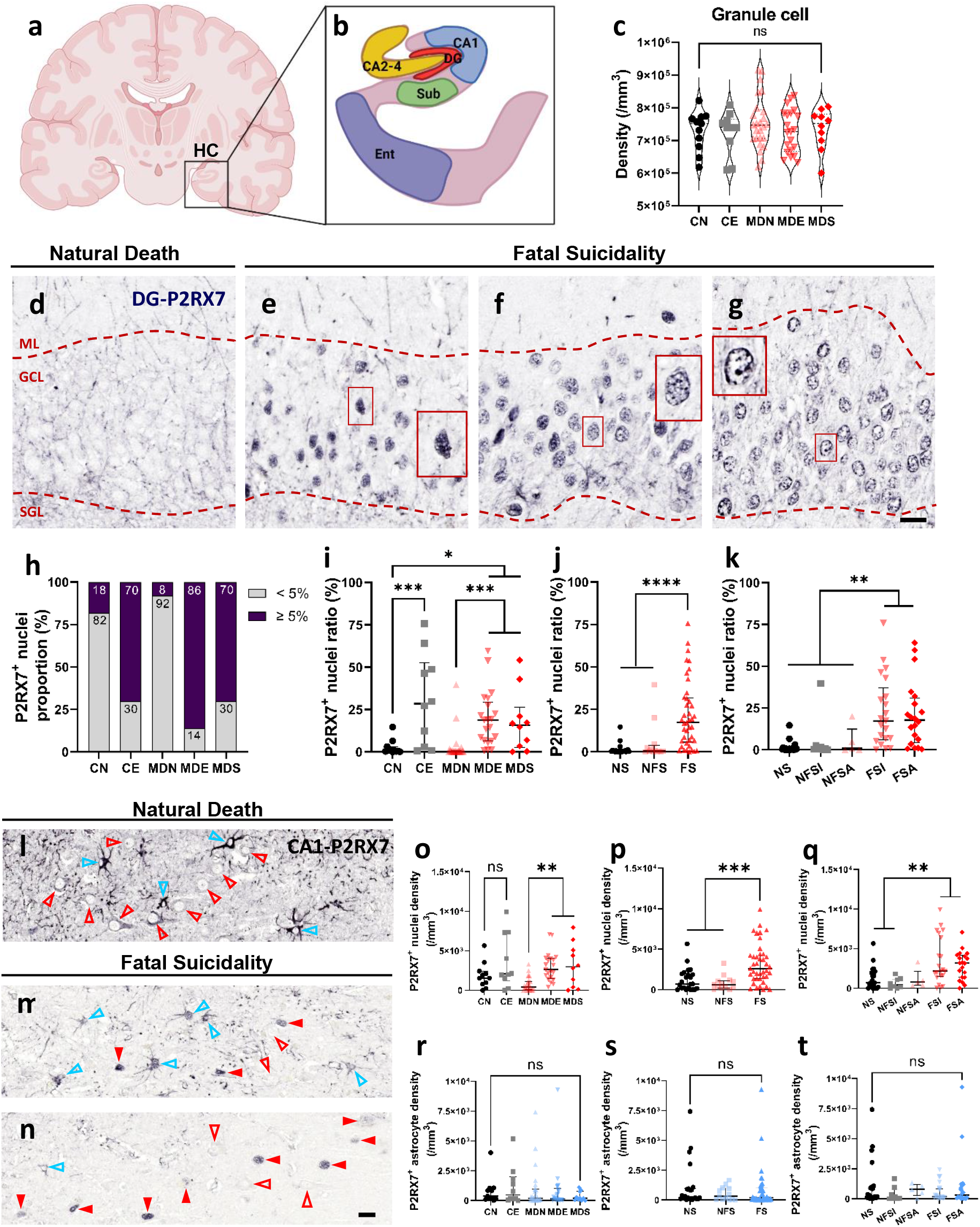
P2RX7^+^ nuclei in the hippocampal formation code for fatal suicidality. Anatomical scheme (**a**) of the hippocampal subregions (**b**) employed in this study. **c** Granule cell density in the DG of different groups is stable. **d-k** P2RX7 expression in the DG of individuals without (**d**) and with fatal suicidality (**e**-**g**). Notably, **e-g** present a progressively increased number of P2RX7^+^ nuclei accompanied by elevated chromatin condensation and marginalization. The boxes magnify the positive staining patterns of P2RX7^+^ nuclei in the GCL. **h** shows higher proportions of individuals with over 5% granule cells that express nuclear P2RX7, and relatively more granular nuclei express P2RX7 in both the Ctr and MD groups in relation to fatal suicidality (**i**). **j** and **k** show a dramatic increase of P2RX7^+^ nuclei in subjects who had fatal suicidality, compared to no or non-fatal suicidality. **l-t** P2RX7 expression in the CA1 of a control subject (**l**) and two depressed patients with fatal suicidality (**m** and **n**). Red solid arrowheads point to neurons with P2RX7^+^ nuclei. Red empty arrowheads point to neurons without nuclear P2RX7 staining. Blue empty arrowheads point to P2RX7^+^ astrocytes. **o** shows the increases of P2RX7^+^ nuclei in the CA1 in MD patients with fatal suicidality, but there is no difference between non-psychiatric control subsets. **p** and **q** reveal dramatic increases of P2RX7^+^ nuclei in subjects with fatal suicidality compared to no or non-fatal suicidality. **r-t** P2RX7-expressing astrocytes in the CA1 did not show differences in relation to suicide or MD. Abbreviations: CA, *cornu ammonis;* CE, control subjects who died by legal euthanasia; CN, control subjects who died by natural causes; DG, dentate gyrus; Ent, entorhinal cortex; FS, fatal suicidality; FSA, suicide attempters who died by suicide or legal euthanasia; FSI, suicide ideators who died by suicide or legal euthanasia; GCL, granule cell layer; HC, hippocampus; MDE, individuals with MD who died by legal euthanasia; MDN, individuals with MD who died by natural causes; MDS, individuals with MD who died by suicide; ML, **molecular layer;** NFS, non-fatal suicidality; NFSA, non-fatal suicide attempts; NFSI, non-fatal suicidal ideations; NS, no suicidality; ns, no significance; **SGL, subgranular layer;** Sub, subiculum. Scale bar, 15 μm. Data are presented as medians with interquartile ranges. * indicates 0.01 ⩽*p* < 0.05, ** indicates 0.001 ⩽*p* < 0.01, *** indicates 0.0001 ⩽*p* < 0.001, **** indicates*p* < 0.0001.

The samples were re-stratified according to the suicidal classification and consequence as mentioned above. We found that in subjects with fatal suicidality, a much higher ratio of P2RX7^+^ nuclei was present than in subjects with a history of non-fatal suicidality or subjects without an inclination for suicide (FS vs NFS: *p* < 0.0001; FS vs NS: *p* < 0.0001; **Fig. 1j**). The ratio of P2RX7^+^ nuclei in the group of non-fatal suicidality subjects was not different from the ratio in non-suicidal subjects (NFS vs NS: *p* = 0.77). Subsequently, fatal and non-fatal suicidalities were subdivided into suicidal ideations and suicide attempts to see whether the suicidal outcome might affect these changes. Relative to non-fatal suicidal ideations, subjects with fatal suicidal ideations showed a significant increase of P2RX7^+^ nuclei compared to non-suicidal cases (FSI vs NFSI: *p* = 0.0007; **Fig. 1k**). Similarly, increased ratios were found in fatal suicidal attempters compared to non-fatal suicidal attempters (FSA vs NFSA: *p* = 0.007; **Fig. 1k**). Subjects with suicidal ideations or with a history of suicide attempt(s) did not show differences within the same prognosis (NFSA vs NFSI: *p* = 0.87; FSA vs FSI: *p* = 0.96; **Fig. 1k**). In addition, when individuals with fatal suicide attempts were subdivided into single attempts and repeated attempts, no differences were found between these groups (*p* = 0.31). Similar changes were observed for P2RX7^+^ nuclei densities (**Fig. S1a-c**).

In contrast to the DG, no differences were present in the CA1 of the controls who died by natural causes as compared to legal euthanasia (CN vs CE: *p* = 0.28; **Fig. 1o**). P2RX7^+^ nuclei in the CA2-4 presented increased density in individuals who died by legal euthanasia compared to natural causes (**Fig. S1j**). In the Sub and Ent, such differences were found only within MD subsets (**Fig. S2d** and **S2m**). Apart from the P2RX7^+^ nuclei, we quantified P2RX7^+^ astrocytes in the hippocampal subregions (CA1 and Sub), but we did not find differences in relation to suicide or MD (**Fig. 1r-t**). Detailed information about the quantification of P2RX7^+^ nuclei in the hippocampal subareas is presented in <u>Supplementary **Fig. S1**-**S2**</u>. The total expression of P2RX7 and GFAP in the hippocampal subfields did not show differences (**Fig. S3** and **S4**).

### Microglial priming in the hippocampal formation correlates with suicidal consequences in mood disorders

Suicide is strongly linked to disrupted microglial homeostasis and phagocytosis in the human brain. To estimate the phagocytotic capacity versus microglial homeostasis, we determined the ratio of the phagosome-to-soma IOD CD68/P2RY12. This ratio was generally decreased throughout hippocampal subregions of MD patients as compared to the controls (**Fig. 2a-e**). Subset analysis showed that, except for the Sub, these reduced ratios in the other subfields mainly resulted from depressed patients with fatal suicidality (**Fig. 2f-j**). Since their average age was younger than non-suicidal controls (**Table S1**), we split the data into middle age and old age (below or above 70 years old) and compared them separately with the controls who died of natural causes. The significant ratio reductions persisted, except for the Sub (**Fig. S5**).

**Figure 2.**
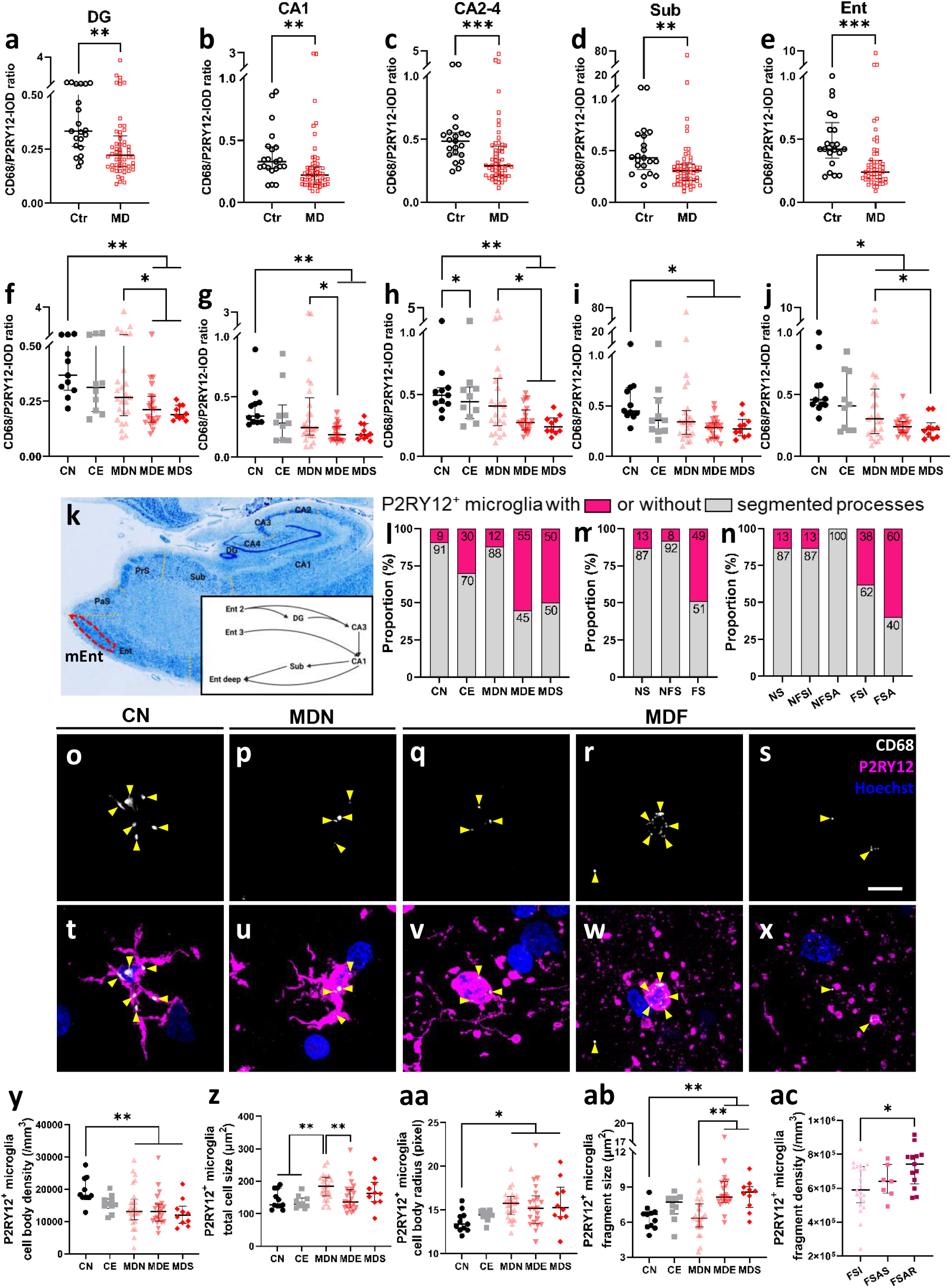
Microglial priming in the hippocampus reflects the suicidal process in mood disorders. **a-j** Quantitative analysis of the ratio of CD68 to P2RY12 shows an overall decline throughout the hippocampal subregions of MD patients (**a-e**), which is mainly due to the reduction in depressed patients who had fatal suicidality (**f-j**). **k** Anatomical localization of the mEnt (superficial layers) following thionine staining. The hippocampal tri-synaptic circuit is schematically miniaturized in the frame. **l-n** display the distribution of deformed P2RY12^+^ microglia in different subgroups in relation to MD and suicidality. Confocal images of CD68 and P2RY12 immunostaining in the mEnt exhibit diminished CD68 expression in MD (**p**-**s**) versus Ctr (**o**). Co-labeled P2RY12^+^ microglia show typical morphological phenotypes from ramified microglia (**t**) in CN, reactive microglia characterized by enlarged cell bodies and bushy processes (**u**) in the MDN, swelling microglia with segmented processes (**v**) or with disconnected microspheres and debris (**w**), and microspheres and debris without microglial cell bodies (**x**) in MD patients with fatal suicidality. Yellow solid arrowheads point to CD68^+^ phagosomes. Imaging analyses show an overall decline of microglial density in the mEnt of patients with depression (**y**). **z** By quantifying the size of microglia, there is an increase in the MDN relative to the controls, and a reduction in the MDE compared to the MDN. **aa** shows an overall increase of P2RY12^+^ microglial cell body radius in MD versus CN. **ab** reveals that the size of microglial fragments is larger in depressed patients with fatal suicidality in comparison to the individuals who died naturally. **ac** In individuals with fatal suicidality, the density of microglial fragments is significantly higher in repeated suicide attempters than in suicidal ideators. Abbreviations: FSAR, FS with repeated suicide attempts; FSAS, FS with a single suicide attempt; MDF, MD patients with fatal suicidality; mEnt, medial entorhinal cortex; PaS, parasubiculum; PrS, presubiculum. Scale bar, **s** 15 μm. Data are presented as medians with interquartile ranges. * indicates 0.01 ⩽*p* < 0.05, ** indicates 0.001 ⩽*p* < 0.01, *** indicates 0.0001 ⩽*p* < 0.001.

Surprisingly, approximately half of the individuals with fatal suicidality showed in the superficial layers of the medial Ent (mEnt) widely distributed P2RY12^+^ microglia with segmented processes and microspheres (**Fig. 2k-m** and **Fig. S6a-c**). Subgroup analysis showed that these heteromorphic microglia were more often detected in fatal suicide attempters than in fatal suicidal ideators (*p* = 0.002, **Fig. 2n**). The morphological distinctions between ramified microglia, reactive microglia, and swollen microglia with segmented processes were confirmed by 3D immunofluorescence reconstructions (**Fig. 2o-x, S6d-g**).

We subsequently quantified the P2RY12^+^ microglia in the mEnt and found their numbers to be reduced in MD compared to CN (*p* = 0.004, **Fig. 2y**). However, the size of the microglia was larger in MDN than in the other subgroups (*p* = 0.003, **Fig. 2z**). To further estimate the extent of cellular deformations, we measured the radius of microglial cell bodies and found longer radii in MD than in CN (*p* = 0.01, **Fig. 2aa**). We also determined the size of microglial fragments and found that they were larger in depressed patients who had fatal suicidality than in subjects with natural death (*p* < 0.0001, **Fig. 2ab**). Notably, among individuals with fatal suicidality, the density of microglial fragments was higher in people who attempted suicide repeatedly than in those who only had suicidal thoughts (*p* = 0.01, **Fig. 2ac**). These malformed microglia were absent in the other hippocampal subareas of the same cohort. Detailed information referring to the quantification is provided in <u>Supplementary **Fig. S6**</u>.

### Selective activation of purinergic glia in the supraoptic and paraventricular nuclei in suicide and mood disorders

The volumes of selected hypothalamic nuclei were determined based on the thionine-delineated magnocellular neurons in the SON, and on both magnocellular and parvocellular neurons in the PVN. Surprisingly, in MD, individuals who died of legal euthanasia or completed suicide exhibited decreases in the SON volumes of 24.8% and 39.3%, respectively, compared to those who died of natural causes (MDE vs MDN: *p* = 0.028; MDS vs MDN: *p* = 0.005, **Fig. 3c**). By contrast, the PVN volume between the five groups was not different (*p* = 0.72, **Fig. 3d**). The volume of the PVN and SON showed no difference between patients with MD and the controls (PVN: *p* = 0.64; SON: *p* = 0.97, **Fig. S7a** and **S7b**).

**Figure 3.**
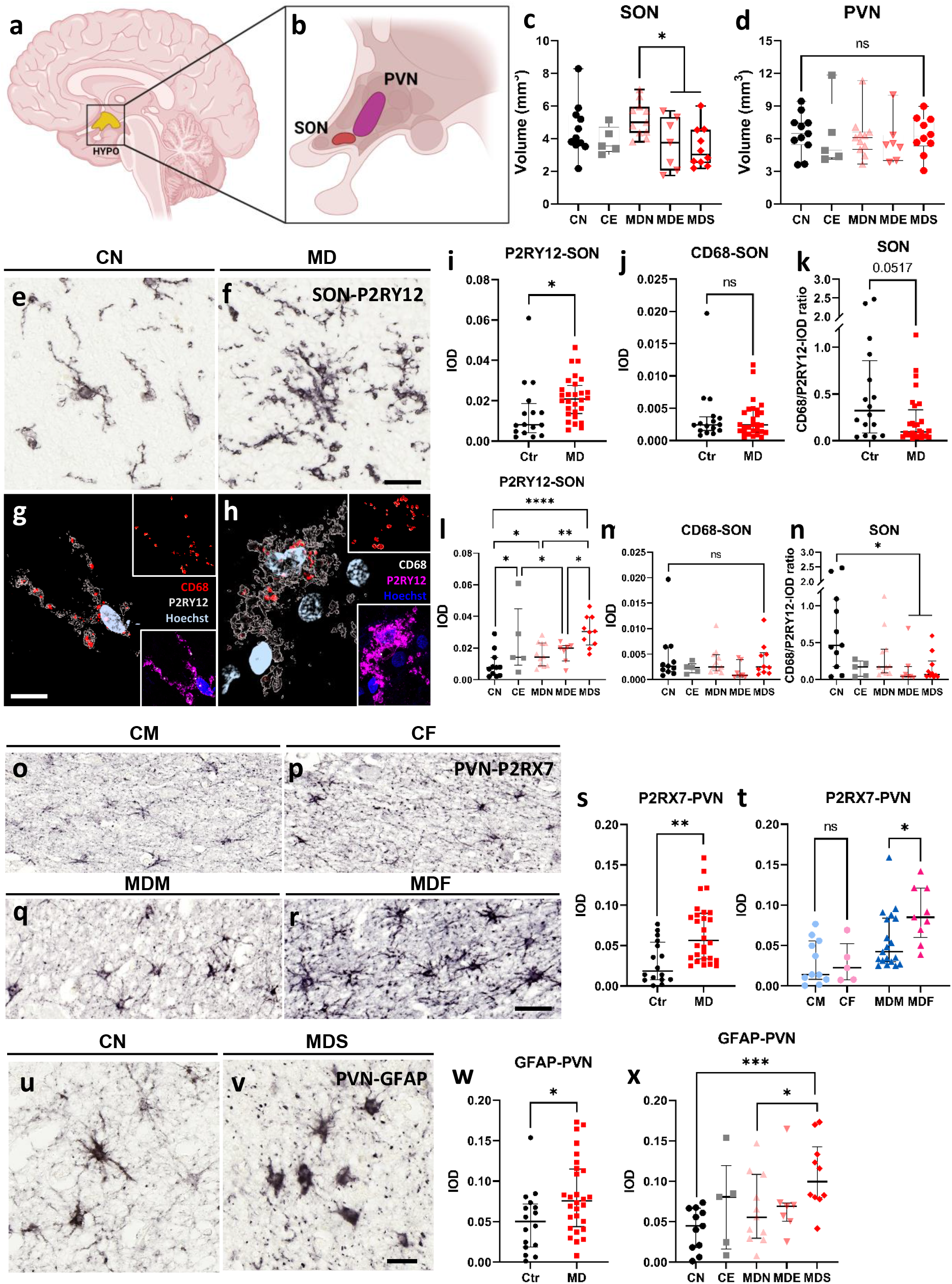
Activated purinergic glia in hypothalamic nuclei are specific to suicide or mood disorders. The anatomical scheme (**a**) of the hypothalamic nuclei studied (**b**). In MD, patients with fatal suicidality show a volumetric shrinkage in the SON (**c**), but not PVN (**d**). **e-h** P2RY12^+^ microglia in the SON of a control subject (**e**) and an MD patient (**f**). Spatial reconstruction specifies CD68^+^ phagosomes in microglia of a non-suicidal control (**g**) and a depressed patient who died of suicide (**h**). Images miniaturized in the top right reveal the reconstructed surfaces of CD68^+^ phagosomes. Images miniaturized in the bottom right show their raw morphology. **i-n** The quantitative analysis in relation to MD, suicidality, and completed suicide. **o-r** display the increase of paraventricular P2RX7 expression in MD (**s**) with a gender preference for females over males, which is absent among the controls (**t**). **o** CM, **p** CF, **q** MDM, **r** MDF. **u** and **v** show GFAP^+^ astrocytes in the PVN of a control subject (quiescent, **u**) and a depressed patient who completed suicide (reactive, **v**). **w** and **x** are the quantitative analyses that demonstrate a specific increase in GFAP expression in MD, which is largely due to accomplished suicide compared to subjects who did not have suicidality. Abbreviations: HYPO, hypothalamus; CF, non-psychiatric females; CM, non-psychiatric males; MDF, female patients with mood disorders; MDM, male patients with mood disorders; PVN, paraventricular nucleus; SON, supraoptic nucleus. Scale bar, 20 μm, except for 10 μm in **Fig. 3g**. Data are presented as medians with interquartile ranges. * indicates 0.01 ⩽*p* < 0.05, ** indicates 0.001 ⩽*p* < 0.01, *** indicates 0.0001 ⩽*p* < 0.001, **** indicates*p* < 0.0001.

Increases of P2RY12 expression were found in the SON and PVN of MD patients relative to the controls (SON: ratio = 2.59, *p* = 0.03, PVN: ratio = 1.41, *p* = 0.049, **Fig. 3e, 3f, 3i** and **S7c**). The microscopy of these P2RY12^+^ microglia revealed increases in size, thickened and bushy processes with retractions. In addition to MD, a higher expression of P2RY12^+^ microglia was present in the SON of subjects with fatal suicidality, which was absent in the PVN (**Fig. 3l** and **S7f**). We further incorporated CD68 expression and found a decreasing trend of CD68/P2RY12 IOD ratio in MD (*p* = 0.052, **Fig. 3k**), which was mainly contributed by subjects who had fatal suicidality compared to the controls without suicidal intension (**Fig. 3n**). However, the expression of CD68 in the hypothalamic nuclei did not change in relation to suicide or depression (**Fig. 3j** and **3m**).

In the PVN of depressed patients, P2RX7 showed over three times higher expression than the controls (*p* = 0.008, **Fig. 3o-s**). In subset analyses, we found female MD patients had two times higher P2RX7 expression in the PVN than their male counterparts (MDF vs MDM: *p* = 0.02, **Fig. 3t**). This sex difference was not observed in the controls (CF vs CM: *p* = 0.78). P2RX7 expression in the SON did not show changes (**Fig. S7i** and **S7j**).

GFAP-positive astrocytes also presented higher expression in the PVN of depression (*p* = 0.049, **Fig. 3u-w**). Elevated expression was found in subjects who died by suicide compared to those who died of natural causes (MDS vs MDN: ratio = 1.79, *p* = 0.01; MDS vs CN: ratio = 2.22, *p* = 0.0002, **Fig. 3x**). However, no differences were shown due to the presence of MD or suicidal ideation. By contrast, GFAP in the SON did not show changes (**Fig. S7k** and **S7l**). For more detailed glial expression in the hypothalamus see <u>Supplementary **Fig. S7**</u>.

### P2RX7 expression decreased in the prefrontal cortex in suicide and major depressive disorder

The expression of P2RX7 in the PFC (DLPFC and ACC) of MD subjects was determined at the transcriptional level. MDD, but not BD patients, had lower prefrontal P2RX7 expression compared to the controls (DLPFC: MDD vs Ctr: fold change = −1.24, *p* = 0.002, MDD vs BD: fold change = −1.29, *p* = 0.03, BD vs Ctr: *p* = 0.41; ACC: MDD vs Ctr: fold change = −1.81,*p* < 0.0001, MDD vs BD: fold change = −1.81,*p* < 0.0001, BD vs Ctr: *p* = 0.78; **Fig. 4b** and **4c**). In addition to completed suicide, the possible impact of psychotic features, which are linked to a clinically high risk for suicide attempts, was examined by subset analyses. Decreased expression of P2RX7 appeared to be more related to suicide in the DLPFC than in the ACC (DLPFC: MDDNS vs Ctr: *p* = 0.14; ACC: MDDNS vs Ctr: fold change = −1.85,*p* < 0.0001; **Fig. 4d** and **4e**). The presence of psychotic features did not affect the P2RX7 expression in the MDD cohort, including the DLPFC and ACC (**Fig. 4f** and **4g**). We did not find significant alterations in P2RX7 transcripts related to suicide or psychotic features in BD (**Fig. 4h-k**).

**Figure 4.**
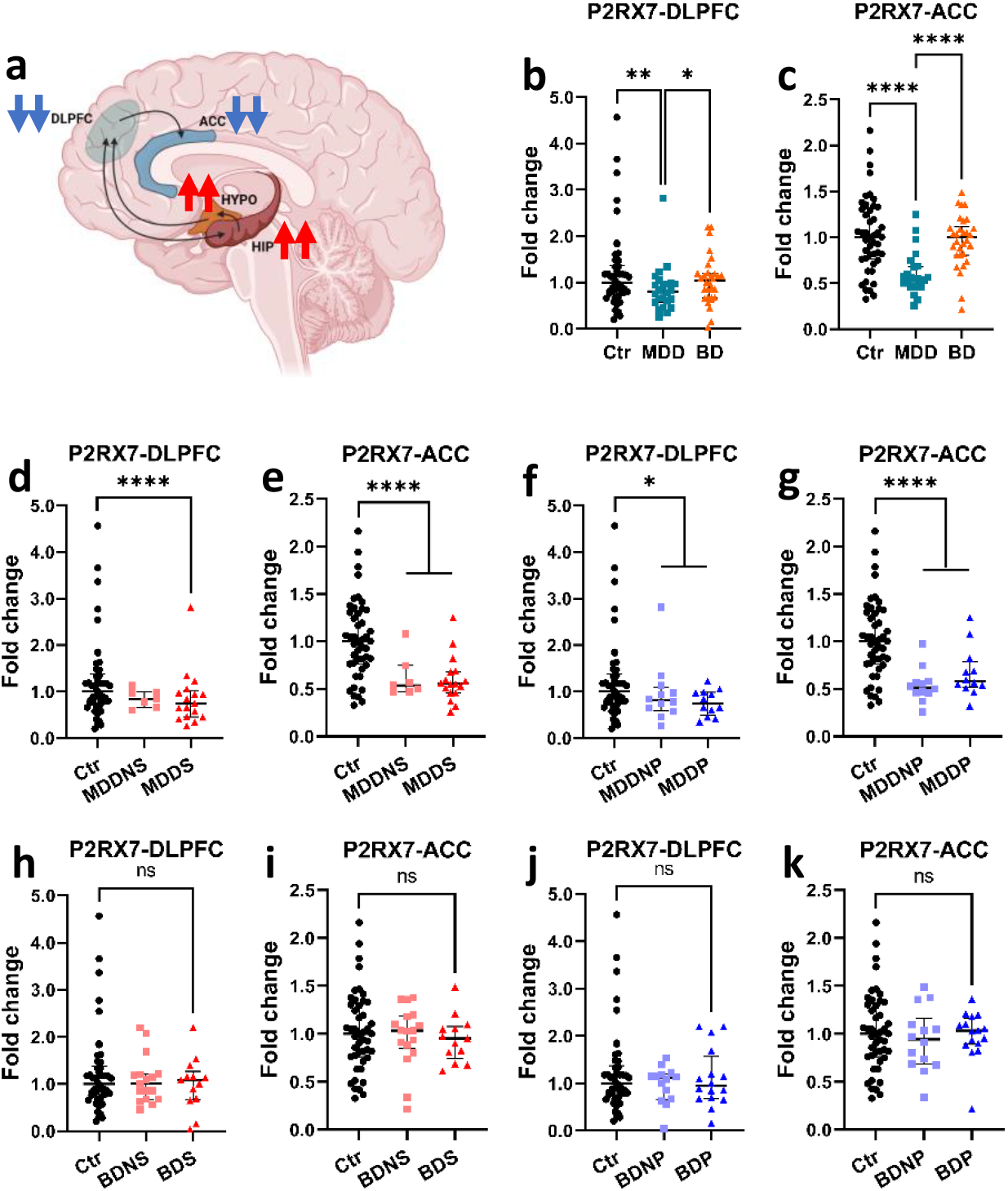
Diminished P2RX7 transcripts in the dorsolateral prefrontal cortex of major depressive disorder are associated with completed suicide. **a** Anatomical scheme of the DLPFC and ACC, and the unified neurocircuitry map of stress-associated purinergic receptors in suicide. Blue/red arrows indicate reduced/elevated purinergic receptor expression, respectively. **b** and **c** show that P2RX7 expression in the PFC is reduced in MDD but not in BD. In the DLPFC of MDD, this reduction is related to completed suicide (**d**), while in the ACC, both MDD and completed suicide present declined P2RX7 expression (**e**). **f** and **g** show that psychotic features do not confound the results. **h-k** shows that the P2RX7 transcripts in the PFC of BD are not altered in completed suicide or psychotic features. Abbreviations: ACC, anterior cingulate cortex; BD, bipolar disorder; BDNP, **bipolar disorder without psychotic features;** BDNS, **bipolar disorder without suicide; BDP, bipolar disorder with psychotic features; BDS, bipolar disorder with suicide**; Ctr, non-psychiatric controls; DLPFC, dorsolateral prefrontal cortex; MDD, major depressive disorder; MDDNP, **major depressive disorder without psychotic features;** MDDNS, **major depressive disorder without suicide;** MDDP, **major depressive disorder with psychotic features;** MDDS, **major depressive disorder with suicide; PFC, prefrontal cortex.** Data are presented as medians with interquartile ranges. * indicates 0.01 ⩽*p* < 0.05, ** indicates 0.001 ⩽*p* < 0.01, **** indicates*p* < 0.0001.

### Putative confounders

Confounder analysis was performed for age, sex, postmortem delay (PMD), clock time of death, and brain pH. Significantly, the presence of P2RX7^+^ nuclei sharply dropped with prolonged PMDs in the hippocampal subfields of donors with fatal suicidality, which was barely detectable 20 hours after death (**Fig. S8a-e**). These negative correlations were absent in donors who died of natural causes (**Fig. S8f-o**). We investigated whether this negative correlation could be alleviated by excluding the intervals over 12h, 9h, and 6h, separately. However, in the DG, P2RX7^+^ nuclei diminished by the PMDs remained evident, as the share of P2RX7^+^ nuclei in the granule cells of fatal suicidality increased more within shorter postmortem intervals (**Fig. S8p-r**). In addition, CD68/P2RY12 ratios throughout the hippocampus of subjects with fatal suicidality were found to increase with age (**Fig. S9a-e**). In individuals who had temporary or transient suicide tendencies, CD68 expression in the CA2-4, Sub and Ent positively correlated with age, with the Ent showing the highest significance (**Fig. S9f-j**). These age-related correlations did not affect our conclusions.

For the PFC, we examined the parameters used to match observations over groups and found that in BD subjects, P2RX7 expression was positively correlated with age in the ACC (Rho = 0.497, *p* = 0.006; **Fig. S9k**), which seemed to be explained by patients with psychotic features who died naturally (Rho = 0.670, *p* = 0.006; **Fig. S9l-o**). In addition, there was a positive correlation between P2RX7 mRNA and lifetime fluphenazine equivalents intake in the ACC of individuals with MD (Rho = 0.594, *p* = 0.0004), which was absent in the DLPFC.

## Discussion

Biological indicators grounded in key brain regions distinguish suicide neuropathology from comorbid psychiatric disorders. Our current findings revealed four possibilities concerning the differences and similarities between suicide and MD. (1) Fatal suicidality has distinct molecular characteristics, which are independent of the psychiatric background. In this study, it was typically shown by the granule cell nuclei in the DG that expressed P2RX7; (2) When a neuropsychiatric disorder is comorbid, suicide/suicidality-associated molecular alterations occur as symbols of disease progression. For instance, we observed nuclear P2RX7 expression in the CA1 and P2RY12^+^ microglia with segmented processes in the mEnt of MD patients with fatal suicidality; (3) Molecules are sensitive to both suicide and psychiatric disorders, for example, the P2RY12 expression in the SON; (4) Completed suicide or the development of suicidality do not confound molecular changes related to pre-existing psychiatric disorders, as shown by the expression of P2RX7 and P2RY12 in the PVN.

In this study, the most important finding is the extensive and ultra-high presence of P2RX7^+^ nuclei in the GCL of individuals who had intense and persistent suicidal ideations. This striking rise appeared not only in MD patients but also in non-psychiatric controls who had a record of fatal suicidality (i.e., individuals who died by legal euthanasia), indicating that suicide might have a profile of genes homogenously rooted in non-psychiatric populations as well as psychiatric patients. By targeting these key genes and brain regions involved in suicide, therapies may be designed for suicidal individuals who are unable of receiving mental care.

Psychological stress is sensed by the innate immune system in the brain via the ATP/P2RX7 pathway ^21^. Aberrant P2RX7 expression in neural cells dysregulates neuronal-glial homeostasis and cognition, which may trigger the emergence of depressive symptoms ^14,22–26^. The DNA promoter of the *P2RX7* gene has a series of binding sites for nuclear transcription factors that biologically adapt and modify gene expression ^27^. Animal studies have demonstrated that guanosine administration, through its binding site on *P2RX7*, can inhibit ATP activity and display antidepressant and pro-neurogenic effects in the hippocampus ^27–29^. In the plasma of treatment-resistant depressed patients, a subpopulation at a high risk of suicide, P2RX7 transcripts were found to correlate with the glucocorticoid receptor, a key stress-responsive transcription factor ^30^. We suggest that the P2RX7 expression in the granule cell nuclei may provide a target for the interactions between neuroinflammation and glucocorticoids ^(31,32^. In addition, chromatin condensation and marginalization in the dentate P2RX7^+^ nuclei may be pathological features of neurodegeneration. P2RX7 initiates programmed cell death by activating caspases ^33^. Given that the overall density of granule cells in the DG remains unchanged, future research should focus on the earlier stages of neuronal apoptosis in the DG of fatal suicidal cases.

Of note, there were no variations in the dentate P2RX7^+^ nuclei between the two fatal suicidal phenotypes (i.e., ideations or attempts). For years, the development of suicidal thoughts was believed to indicate an aggravation of the pre-existing psychiatric disorder ^34^. In our study, however, legal euthanasia donors who did not have diagnosable mental illnesses, but requested euthanasia due to their malignant diseases, also presented a remarkable proportion of P2RX7^+^ nuclei in the DG. Nine out of ten of these donors had specified suicidal ideations but not behaviors. The only attempter in this subset did not contribute an extreme value of P2RX7^+^ nuclei. It has been proven that the heterogeneity of the *P2RX7* gene was strongly linked to cognitive deficits, a key factor in the suicide processes ^35,36^. Therefore, the distinctive presence of P2RX7^+^ nuclei in the DG may provide a pointcut for deciphering the suicide-specific cognitive inflexibility.

An unavoidable question is whether the high-proportioned P2RX7^+^ nuclei in granule cells of fatal suicidal cases serve as a surrogate for one particular cell group or a proxy for several subdivisions that respond to lethal suicidal thoughts. After all, nuclear transcriptional factors and mechanistic insights implicated in these suicide-specific nuclei remain unknown. Assuming neurological substrates of the DG form a molecular fingerprint of fatal suicidality, the question is whether any P2RX7^+^ nuclei subset(s) could distinguish suicidal phenotypes or consequences. As the presence of P2RX7^+^ nuclei rapidly decreased with the extension of the PMD, technological approaches that profile transcriptomic alterations at single-cell resolution may provide a feasible alternative for overcoming PMD-dependent protein degradation in postmortem human DG ^37^.

Microglial homeostasis in human tissue can be mimicked by biomarkers that monitor phagocytic capacity and morphological indices. In this study, the ratio of CD68 to P2RY12, used to reflect the relation between microglial phagocytosis and homeostasis, generally declined in the hippocampus of MD patients ^38^. Subset analysis showed that this dramatic decrease was more associated with fatal suicidality. Additionally, lesioned processes in entorhinal microglia were implicated in suicidal classifications and severity. First, microglial cell bodies in the mEnt of the entire MD cohort that showed reductions in density together with soma enlargement suggested microglial degeneration. Second, the total size of microglia was larger only in depressed patients who died from non-suicidal causes compared to other subsets, while microglial fragments were smaller in these patients than in those with fatal suicidality. This suggests that a significant number of microglial processes were disconnected from cell bodies. Third, the density of these fragments in the fatal suicidality cohort was much higher in repeated suicide attempters than in suicidal ideators, indicating that entorhinal microglial lesions reflect the repetition of suicidal behaviors. Similar segmented processes in microglia have been found in a marmoset model of autism, a neurological disorder with suicide as a leading cause of premature death ^39,40^. In clinical practice, cellular deformability was reported in peripheral blood monocytes, a myelomonocytic relative of microglia, in patients with persistent MDD, in whom the rate of lifetime suicide attempts is over three times higher than in non-chronic MDD ^41,42^ Taken together, these data suggest that morphological features of immune cells can reflect the severity of psychiatric illnesses. The heteromorphic microglia that mark alterations in energy metabolism and immune homeostasis are extensively distributed in the superficial layers of the Ent (Ent2 and Ent3), which contain the major stimulating afferents to the DG that effectuate the impact of antidepressants ^43,44^ Our next step is to determine certain groups of entorhinal neurons that associate with suicidal severity and explore their functional connectivity within the Ent-DG circuitry.

So far, neuroimaging strategies cannot precisely identify the hypothalamic nuclei because their anatomical boundaries lack contrast. We discovered that MD patients with fatal suicidality had reduced SON volumes compared to non-suicidal MD patients. The SON is generally homogeneous and primarily composed of vasopressin neurons, which have been linked to enhanced suicide risk ^45^. However, this activation probably lacks effective regulation of microglial phagocytosis, as CD68 expression was unchanged. We assume that the entire process of suicide may drive persistent hyperactivity in the hypothalamic-pituitary-adrenal axis and the consequent strengthening of negative cortisol feedback to the SON, which may reduce its volume ^46,47^ In the PVN, both P2RX7 and P2RY12 were highly expressed in depressed subjects, indicating a significant extracellular release of ATP and ADP. When compared to non-suicidal individuals, cases of completed suicide had an additionally increased GFAP expression in the PVN, suggesting that hypothalamic astrocytes may have a distinct role in suicide pathophysiology.

In addition, we observed a sex-specific increase of P2RX7 expression in the PVN of female depressed patients. Genetic associations between *P2RX7* mutations and the higher risk of BD occurrence have been reported in females but not males ^48^. In animal stress models, a higher P2RX7 expression in the PVN may likely induce a gender-biased elevation of FosB, a transcription factor that modulates several target genes implicated in depression ^49–51^. Again, sex may be a mechanism-modifying variable with important treatment implications.

In contrast to the subcortical purinergic receptors that were overactivated in suicide, P2RX7 transcripts in the PFC of MDD subjects, but not BD subjects, showed sharp declines when compared to the controls. In MDD, this decrease in the DLPFC may have been caused by P2RX7 depletion only in cases of suicide, which is consistent with other studies ^(7,12^. There were no differences in the ACC between patients with or without actual suicides. Neither did prefrontal P2RX7 expression in BD change in relation to suicide. However, P2RY12 transcripts in the DLPFC of BD donors, as reported in our earlier findings, were lower in non-suicidal patients than in suicide victims and non-psychiatric controls ^5^. Since the PFC is involved in negative feedback regulation of the hypothalamus-pituitary-adrenal axis and hippocampal responses to stress ^52,53^, the findings above suggest that purinergic hypoactivity in the PFC together with hyperactivity in the hippocampus and hypothalamus could reflect a diminished inhibitory connection between cortical and subcortical components. Further, recent studies have found that a *P2RX7* variant plays a role in the vulnerability to develop psychotic features, which have a markedly increased likelihood of subsequent suicide death ^54^. Our results, however, did not provide evidence that this observation could be related to the prefrontal regions.

### Limitations

There are some limitations to this study. First, the detectable number of P2RX7^+^ nuclei in the hippocampus of fatal suicidality subjects significantly dropped over PMD. This has weakened the analysis of possible impacts, such as suicidal classifications, consequences, or reasons for legal euthanasia (physical versus psychiatric diseases). The PMDs among suicide completers were relatively long and recorded by estimation. Practically speaking, some critical molecules may have been degraded, which confines the brain sample inclusion for further research. To retain these suicide biomarkers, we introduced brain samples from donors who died by legal euthanasia and emphasized the imperative role of their short and precise PMDs. Second, P2RX7^+^ nuclei were densely packed in the DG but loosely distributed in the other hippocampal subregions, which resulted in increased data variation and decreased statistical significance. Therefore, we could not determine whether regional specificity or methodological constraints were to blame for the lack of significance between fatal and non-fatal suicidality in the Sub or Ent (**Fig. S2d** and **S2m**). These two hurdles can be technically overcome by follow-up studies using single-nucleus transcriptomics. Third, PFC samples were obtained from the SMRI, from which legal euthanasia cases were not available. We do not have information on prefrontal alterations in relation to fatal suicidal thoughts or attempts. Fourth, the average age of the prefrontal collection was thirty years younger than the other brain cohorts. However, our recent proteomic studies have reported sharply diminished energy metabolism in the PFCs of elderly depressed patients in the NBB cohort, which supported our conclusions ^55^.

## Conclusions

The biological hallmarks of fatal suicidality are molecule-, cell type- and region-specific. They may be either independent or (partially) overlapping with the changes observed in the comorbid psychiatric disorders. We investigated a network linked to suicidality, which was mediated by a mini neurocircuit in the hippocampus, and a major circuit involving the hypothalamus and PFC. Our discovery of a specific group of hippocampal granule cells exclusively associated with fatal suicidality indicates that the DG may contain cellular and molecular signatures for intense and determined suicidal ideations ^35^. Importantly, this suicide-specific feature sharply dropped over PMD, even within our very short intervals. This phenomenon urged the application of brain samples from donors who died by legal euthanasia, to enable the study of suicide biosignatures that were insignificant in previous research. Microglial dyshomeostasis and morphological traits in the mEnt of depression cases were found to be related to the suicidal categories. Similar findings were presented in the hypothalamic SON. Our results suggest that, in individuals with intense and determined suicidality, certain subcortical regions may exhibit increased energy metabolism, which may be due to the deficiency of prefrontal inhibition. Overall, unraveling the molecular basis that identifies suicide can equally provide new therapeutic targets for suicidal individuals with or without mental care.

## Contributors

L. Zhang and D.F. Swaab designed the research project. L. Zhang undertook NBB sample (the hippocampus and hypothalamus) collection and processing, immunohistochemical and immunofluorescent staining, image analysis, and data collection. J. van Heerikhuize designed the quantitative strategies and supervised data processing. R. Balesar and Z. Slabe undertook SMRI sample (the PFC) application and processing, real-time qPCR, and data collection. R.W.H. Verwer and L. Zhang undertook the statistical analysis. F. Corrêa da Silva performed image processing for 3D reconstruction. D.F. Swaab took care of financing the project. L. Zhang wrote the first draft. D.F. Swaab and P.J. Lucassen amended the manuscript. All authors have approved and contributed to the final manuscript.

## Acknowledgments

We thank A.A. Sluiter, C. Yi, M. de Lange, A.R. Muller, X. Chen, H. Tan, and E. Salta for their theoretical and technical support. We appreciate the dedication of everyone from SMRI and NBB.

