## Supplementary information for "Stress-associated purinergic receptors code for fatal suicidality in the hippocampal-hypothalamic-prefrontal circuit"

### Supplementary Materials and Methods

#### Neuropathological admittance and sample matching

**NBB collections**: Patients with MD had been diagnosed by licensed psychiatrists during their lifetime according to the Diagnostic and Statistical Manual of Mental Disorder (DSM)-III, IIIR, and IV. A neuropathologist systematically examined each brain to exclude pathological alterations. Subjects with evidence of neurological disorders or with Braak stage III or higher for Alzheimer's were excluded [1]. Subjects had no significant history of substance dependence 10 years before death. Matching for potentially confounding factors was non-parametrically tested. This included age, sex, post-mortem delay (PMD), month of death, clock time of death (CTD), brain weight, and the pH of cerebrospinal fluid. We noticed that ages of subjects with MD who died of suicide or legal euthanasia were younger than the other subjects. Since legal euthanasia is generally performed during working hours, CTDs of euthanasia groups were, therefore, significantly different from the other groups. In addition, PMDs of suicide completers were estimated values, while they were precise values in the other subsets. For demographic and clinic-pathological information see **Table S1a-d**.

**SMRI collections:** Permission for the use of brain material was provided by the next of kin. Diagnoses were based on the DSM-IV. All brain regions were microscopically examined to exclude patients having pathological signs of neurodegeneration or other lesions. Exclusion criteria included subjects with a history of seizures or other neurological disorders that might affect brain pathology. All groups were matched for age, brain pH, PMD, brain weight, history, and severity of alcohol/substance abuse. For demographic information see **Table S1e-f**.

**Table S1a Demographic information of Suicide-HIPPOCAMPUS collection (NBB)**

| Diagnosis Cause of death | Cases, n | Psychiatric diagnosis (MDD/BD) | Reason for euthanasia (Physical diseases/MD)[1] | Age (y)[2] | Sex (M/F) | PMD (h)[2] | Brain pH[2] | BW (g)[2] |
|---|---|---|---|---|---|---|---|---|
| **MD** | 56 | 37/19 | - | 66.5 (23-93) | 21/35 | 6:58 (3:55-20:35) | 6.62 (5.61-7.30) | 1220 (912-1525) |
| Suicide | 10 | 7/3 | - | 58.5 (43-90) | 2/8 | 9:52 (3:55-20:35) | 6.58 (6.18-6.93) | 1215 (1120-1475) |
| Legal euthanasia | 21 | 18/3 | 9/13[3] | 61 (23-83) | 6/15 | 6:20 (4:10-12:45) | 6.80 (6.48-7.30) | 1210 (1045-1525) |
| Natural causes | 25 | 12/13 | - | 70 (45-93) | 13/12 | 7:00 (4:20-10:20) | 6.50 (5.61-6.95) | 1237 (912-1510) |
| **Ctr** | 21 | - | - | 83 (51-94) | 8/13 | 5:45 (4:15-12:50) | 6.56 (6.05-7.00) | 1259 (1054-1590) |
| Legal euthanasia | 10 | - | 10/0 | 82.5 (51-92) | 2/8 | 5:40 (4:25-7:50) | 6.63 (6.36-6.92) | 1206 (1090-1390) |
| Natural causes | 11 | - | - | 83 (73-94) | 6/5 | 5:50 (4:15-12:50) | 6.49 (6.05-7.00) | 1275 (1054-1590) |

**Abbreviations:** BD, bipolar disorder; BW, brain weight; Ctr, non-psychiatric controls; F, female; g, gram; h, hour; M, male; MD, mood disorders; MDD, major depressive disorder; NBB, Netherlands Brain Bank; PMD, postmortem delay; y, year.

**Note:** 1 In the Netherlands, legal euthanasia will be approved to end unbearable suffering with no prospect of improvement in physical/psychiatric diseases (https://www.government.nl/topics/euthanasia/is-euthanasia-allowed). For the practice of legal euthanasia in the Netherlands see https://www.knmp.nl/downloads/guidelines-for-the-practice-of-euthanasia.pdf.

2 Values are provided with medians (and ranges).

3 One individual who died of legal euthanasia applied on basis of a combination of physical diseases and MD. The others were approved either because of physical diseases or MD.

**Table S1b Clinico-pathological information of patients with suicide, legal euthanasia, and control subjects (Suicide-HIPPOCAMPUS collection, NBB)**

| Sample number | Psychiatric diagnosis | Age (y) | Sex | PMD (h) | CTD | CSF pH | BW (g) | Suicidality | Antipsychotics in last three months | Cause of death |
|---|---|---|---|---|---|---|---|---|---|---|
| MD1 | MDD | 59 | F | 20:35 | 00:00 | 6.53 | 1199 | Repeated attempts | BZD | Suicide |
| MD2 | MDD | 56 | F | 10:30 | 14:00 | 6.42 | 1120 | Repeated attempts | BZD | Suicide, intoxication |
| MD3 | MDD | 62 | F | 11:40 | 16:15 | 6.18 | 1155 | Repeated attempts | BZD | Suicide, strangulation |
| MD4 | BD | 83 | M | 15:25 | 10:15 | 6.5 | 1401 | Single attempt | TeCA | Suicide, strangulation |
| MD5 | MDD | 58 | M | 04:30 | 09:40 | 6.8 | 1470 | Single attempt | TCA, SGA, SSRI | Suicide, intoxication |
| MD6 | BD | 46 | F | 05:45 | 16:45 | N.A. | 1370 | Suicidal ideations | BZD | Suicide, intoxication |
| MD7 | MDD | 85 | F | 04:30 | 12:00 | 6.93 | 1200 | Suicidal ideations | AA, BZD, TCA | Suicide, intoxication |
| MD8 | MDD | 43 | F | 11:20 | 10:00 | N.A. | 1475 | Suicidal ideations | BZD, SSRI | Suicide, hanging |
| MD9 | BD | 55 | F | 09:15 | 09:45 | 6.83 | 1210 | Suicidal ideations | AA, BZD, DPBP, VPA | Suicide, intoxication |
| MD10 | MDD | 90 | F | 03:55 | 11:20 | 6.62 | 1220 | Suicidal ideations | BZD, INN | Suicide, intoxication |
| MD11 | MDD | 60 | F | 04:20 | 16:10 | N.A. | 1074 | Suicidal ideations | Hal | Legal euthanasia |
| MD12 | MDD | 83 | F | 04:05 | 14:25 | 6.75 | 1066 | Single attempt | BZD, TeCA | Legal euthanasia |
| MD13 | MDD | 47 | F | 06:05 | 15:00 | N.A. | 1264 | Repeated attempts | AA, BZD, SARI | Legal euthanasia |
| MD14 | MDD | 73 | F | 05:45 | 15:30 | N.A. | 1205 | Suicidal ideations | TeCA | Legal euthanasia |
| MD15 | MDD | 66 | F | 07:55 | 14:35 | N.A. | 1165 | Repeated attempts | AA, BZD, SNRI | Legal euthanasia |
| MD16 | MDD | 47 | M | 05:55 | 11:35 | 6.67 | 1390 | Repeated attempts | N.A. | Legal euthanasia |
| MD17 | MDD | 66 | F | 10:40 | 17:00 | 6.71 | 1045 | Suicidal ideations | VPA | Legal euthanasia |
| MD18 | MDD | 55 | M | 04:19 | 16:11 | 6.48 | 1190 | Repeated attempts | N.A. | Legal euthanasia |

| | | | | | | | | | | |
|---|---|---|---|---|---|---|---|---|---|---|
| MD19 | MDD | 77 | F | 08:40 | 11:20 | 6.77 | 1120 | Repeated attempts | BZD, INN, nBZD, SSRI | Legal euthanasia |
| MD20 | MDD | 50 | M | 09:40 | 13:00 | 6.8 | 1420 | Repeated attempts | BZD | Legal euthanasia |
| MD21 | MDD | 38 | M | 06:15 | 11:45 | 6.79 | 1525 | Single attempt | BZD | Legal euthanasia |
| MD22 | BD | 45 | F | 07:30 | 13:00 | 6.93 | 1325 | Repeated attempts | SSRI | Legal euthanasia |
| MD23 | MDD | 67 | F | 08:40 | 15:35 | 7.07 | 1205 | Suicidal ideations | BZD | Legal euthanasia |
| MD24 | MDD | 23 | F | 08:35 | 15:45 | N.A. | 1415 | Repeated attempts | AA, BZD, NDRI, TCA | Legal euthanasia |
| MD25 | MDD | 48 | M | 12:45 | 10:15 | 6.76 | 1350 | Single attempt | MAOI | Legal euthanasia |
| MD26 | MDD | 61 | M | 05:45 | 14:05 | N.A. | 1210 | Suicidal ideations | SSRI | Legal euthanasia |
| MD27 | MDD | 74 | F | 06:20 | 13:45 | 6.96 | 1313 | Single attempt | AA, BZD, nBZD, TCA | Legal euthanasia |
| MD28 | BD | 81 | F | 07:35 | 15:25 | 7.3 | 1250 | Repeated attempts | BZD | Legal euthanasia |
| MD29 | BD | 59 | F | 06:35 | 11:35 | 6.92 | 1132 | Single attempt | BZD | Legal euthanasia |
| MD30 | MDD | 72 | F | 04:30 | 19:30 | 6.88 | 1168 | Suicidal ideations | BZD, SSRI, TCA | Legal euthanasia |
| MD31 | MDD | 61 | F | 04:45 | 10:35 | 6.81 | 1290 | Suicidal ideations | BZD, SSRI | Legal euthanasia |
| MD32 | BD | 70 | M | 04:50 | 02:45 | 6.26 | 1442 | No | N.A. | Cardiac arrest |
| MD33 | MDD | 45 | M | 07:00 | 02:30 | 6.55 | 1412 | Suicidal ideations | SSRI | Brain hemorrhage |
| MD34 | MDD | 81 | M | 06:00 | 15:30 | 6.5 | 1280 | No | Hal | Renal insufficiency |
| MD35 | MDD | 70 | M | 07:15 | 08:00 | 6.5 | 1380 | Single attempt | BZD, MAOI, SGA, SNRI, TCA | Respiratory insufficiency |
| MD36 | BD | 80 | F | 09:30 | 09:30 | 6.33 | 1140 | No | SSRI, VPA | Acute cardiac arrest |
| MD37 | MDD | 93 | F | 04:20 | 04:55 | 6.8 | 1023 | Suicidal ideations | BZD, SSRI | Pneumonia |
| MD38 | MDD | 80 | F | 06:40 | 16:30 | 6.62 | 1237 | Suicidal ideations | BZD, TA, TCA | Sudden death |
| MD39 | MDD | 68 | M | 08:55 | 05:45 | 6.82 | 1510 | Repeated attempts | BZD, SSRI | Sudden death |

| | | | | | | | | | | |
|---|---|---|---|---|---|---|---|---|---|---|
| MD40 | BD | 66 | M | 07:35 | 16:40 | 5.83 | 1320 | Repeated attempts | BZD, Li | Acute myeloid leukemia |
| MD41 | BD | 58 | M | 09:15 | 12:40 | N.A. | 1380 | Suicidal ideations | TCA | Atrial fibrillation |
| MD42 | MDD | 69 | M | 04:40 | 20:10 | 6.63 | 1145 | Suicidal ideations | BZD, SSRI | Multiorgan failure |
| MD43 | MDD | 88 | F | 10:00 | 09:00 | 6.26 | 1160 | No | BZD, SSRI | Burns, dehydration |
| MD44 | BD | 70 | M | 06:50 | 21:00 | 6.19 | 1275 | No | BZD, LTG, VPA | Pulmonary infection |
| MD45 | BD | 64 | F | 05:00 | 19:15 | 6.76 | 1170 | No | BZD, TCA | CVA |
| MD46 | MDD | 69 | F | 06:55 | 09:15 | 6.28 | 1470 | No | nBZD, SSRI, TeCA | Congestive heart failure |
| MD47 | MDD | 58 | F | 07:20 | 23:00 | 5.61 | 1295 | Suicidal ideations | BZD, DA stabilizer, SSRI | Renal insufficiency |
| MD48 | BD | 65 | M | 04:50 | 18:30 | N.A. | 1305 | No | BZD, Li, SGA | Metastatic colon carcinoma |
| MD49 | MDD | 86 | F | 07:30 | 08:30 | 6.57 | 912 | No | AA, BZD | Metastatic ovarian cancer |
| MD50 | BD | 62 | M | 07:30 | 09:05 | 6.4 | 1005 | No | AA, BZD | Palliative sedation |
| MD51 | BD | 82 | F | 10:15 | 18:55 | 6.6 | 1220 | Suicidal ideations | AA, BZD, Li, LTG | Myocardial infarction |
| MD52 | BD | 70 | F | 06:00 | 00:35 | 6.95 | 971 | No | BZD | Kidney failure |
| MD53 | BD | 78 | F | 05:45 | 03:30 | 6.5 | 1010 | Suicidal ideations | N.A. | Acute kidney injury |
| MD54 | MDD | 91 | M | 06:10 | 00:30 | 6.42 | 1064 | No | BZD, MAOI, SGA, VPA | Respiratory insufficiency |
| MD55 | BD | 70 | F | 07:00 | 14:10 | 6.37 | 985 | Repeated attempts | BZD, Li, SGA, VPA | Sepsis |
| MD56 | BD | 68 | M | 10:20 | 01:40 | 6.41 | 1430 | No | BZD | CVA |
| **Median** | - | 66.5 | - | 06:58 | - | 6.62 | 1220 | - | - | - |
| Ctr1 | - | 62 | F | 07:55 | 11:15 | 6.4 | 1115 | Suicidal ideations | BZD | Legal euthanasia |
| Ctr2 | - | 51 | F | 05:36 | 17:59 | 6.92 | 1090 | Suicidal ideations | TCA | Legal euthanasia |
| Ctr3 | - | 89 | F | 06:10 | 13:35 | 6.36 | 1199 | Suicidal ideations | - | Legal euthanasia |
| Ctr4 | - | 89 | F | 04:45 | 16:15 | 6.67 | 1259 | Suicidal ideations | TCA | Legal euthanasia |

| | | | | | | | | | | |
|---|---|---|---|---|---|---|---|---|---|---|
| Ctr5 | - | 75 | F | 05:25 | 18:10 | 6.75 | 1320 | Suicidal ideations | BZD | Legal euthanasia |
| Ctr6 | - | 92 | F | 04:25 | 13:25 | 6.65 | 1212 | Repeated attempts | - | Legal euthanasia |
| Ctr7 | - | 92 | F | 07:45 | 10:00 | 6.71 | 1200 | Suicidal ideations | - | Legal euthanasia |
| Ctr8 | - | 65 | M | 05:45 | 18:25 | 6.55 | 1390 | Suicidal ideations | BZD, LEV | Legal euthanasia |
| Ctr9 | - | 91 | M | 05:05 | 10:20 | 6.55 | 1175 | Suicidal ideations | - | Legal euthanasia |
| Ctr10 | - | 76 | F | 06:55 | 16:50 | 6.61 | 1295 | Suicidal ideations | - | Legal euthanasia |
| Ctr11 | - | 82 | M | 12:55 | 05:00 | 6.21 | 1406 | No | - | Sudden death. |
| Ctr12 | - | 83 | M | 05:45 | 03:00 | 6.35 | 1590 | Single attempts | BZD, Hal, SSRI, VPA | Pneumonia |
| Ctr13 | - | 91 | F | 04:15 | 13:20 | 6.5 | 1054 | No | - | Heart infarction |
| Ctr14 | - | 82 | F | 05:15 | 18:55 | 6.34 | 1221 | No | BZD | Thoracic aortic dissection |
| Ctr15 | - | 73 | M | 04:25 | 05:05 | 7 | 1285 | No | BZD, TCA | Pneumonia |
| Ctr16 | - | 82 | F | 06:20 | 08:30 | N.A. | 1275 | No | - | Physical deterioration |
| Ctr17 | - | 94 | M | 05:30 | 16:15 | 6.57 | 1229 | No | BZD | Metastatic lung cancer |
| Ctr18 | - | 87 | M | 06:20 | 13:10 | 6.47 | 1275 | No | BZD | Renal insufficiency |
| Ctr19 | - | 85 | F | 06:50 | 09:20 | 6.84 | 1190 | No | BZD, Li | Metastatic lung cancer |
| Ctr20 | - | 82 | M | 05:55 | 18:45 | 6.05 | 1580 | No | BZD | Prostate cancer |
| Ctr21 | - | 84 | F | 07:50 | 17:55 | 6.59 | 1345 | No | BZD | Colon cancer |
| **Median** | - | 83 | - | 05:45 | - | 6.56 | 1259 | - | - | - |
| ***p*-value** | - | $< 0.0001$ | 0.96 | 0.053 | n.sig | 0.38 | 0.53 | - | - | - |

**Abbreviations:** AA, atypical antidepressant, BD, bipolar disorder; BW, brain weight; BZD, benzodiazepine; CTD, clock time of death; CTR, control; CVA, cardiovascular arrest; DA, dopamine; DPBP, diphenylbutylpiperidine; F, female; g, gram; h, hour; Hal, haloperidol; INN, nemifitide; LEV, levetiracetam; Li, lithium; LTG, lamotrigine; M, male; MAOI, monoamine oxidase inhibitor; MD, mood disorders; MDD, major depressive disorder; N.A., not available; nBZD, nonbenzodiazepines; NDRI, norepinephrine–dopamine reuptake inhibitor; n.sig, non-significant; PMD, postmortem delay; SARI, serotonin antagonist and reuptake inhibitor; SGA, second-generation antipsychotics; SNRI, serotonin-norepinephrine reuptake inhibitor; SSRI, selective serotonin reuptake inhibitor; TA, typical antipsychotics; TCA, tricyclic antidepressants; TeCA, tetracyclic antidepressants; VPA, valproate; y, year.

**Table S1c Demographic information of Suicide-HYPOTHALAMUS collection (NBB)**

| Diagnosis<br>Cause of death | Cases, n | Psychiatric diagnosis (MDD/BD) | Reason for euthanasia (Physical diseases/MD)[1] | Age (y) | Sex (M/F) | PMD (h) | Brain pH | BW (g) |
|---|---|---|---|---|---|---|---|---|
| **MD** | 28 | 22/6 | - | 66 (38-94) | 19/9 | 8:55 (3:30-62:55) | 6.70 (5.61-7.55) | 1291 (985-1670) |
| Suicide | 10 | 9/1 | - | 60 (39-90) | 6/4 | 20:35 (3:55-62:55) | 6.53 (6.18-6.80) | 1300 (1120-1670) |
| Legal euthanasia | 7 | 6/1 | 4/5 | 61 (38-81) | 5/2 | 6:40 (3:30-9:40) | 6.78 (6.70-7.07) | 1283 (1120-1525) |
| Natural causes | 11 | 7/4 | - | 70 (58-94) | 8/3 | 8:55 (4:50-60:20) | 6.37 (5.61-7.55) | 1295 (985-1510) |
| **Ctr** | 16 | - | - | 72 (39-88) | 11/5 | 7:15 (3:20-41:00) | 6.60 (5.37-7.28) | 1259 (1115-1629) |
| Legal euthanasia | 5 | - | 5/0 | 79 (49-83) | 3/2 | 5:45 (3:20-6:30) | 6.80 (6.15-7.28) | 1173 (1121-1364) |
| Natural causes | 11 | - | - | 67 (39-88) | 8/3 | 9:00 (4:35-41:00) | 6.00 (5.37-6.60) | 1383 (1115-1629) |

**Abbreviations:** BD, bipolar disorder; BW, brain weight; Ctr, non-psychiatric controls; F, female; g, gram; h, hour; M, male; MD, mood disorders; MDD, major depressive disorder; NBB, Netherlands Brain Bank; PMD, postmortem delay; y, year.

**Table S1d Clinico-pathological information of patients with suicide, legal euthanasia, and control subjects (Suicide-HYPOTHALAMUS collection, NBB)**

| Sample number | Psychiatric diagnosis | **Age (y)** | **Sex** | **PMD (h)** | **CTD** | **CSF pH** | **BW (g)** | **Suicide attempt** | **Antipsychotics in last three months** | **Cause of death** |
|---|---|---|---|---|---|---|---|---|---|---|
| MD1 | MDD | 50 | M | N.A. | N.A. | N.A. | 1380 | Yes | N.A. | Suicide |
| MD2 | MDD | 39 | M | 41:00 | 00:00 | N.A. | 1670 | Yes | TCA, SSRI | Suicide |
| MD3 | BD | 39 | M | 48:00 | 11:30 | N.A. | 1220 | Yes | BZD, RIMA, SSRI | Suicide, intoxication |
| MD4 | MDD | 79 | M | 21:10 | 17:50 | N.A. | 1530 | Yes | BZD, SSRI | Suicide, high fall |
| MD5 | MDD | 74 | M | 62:55 | 17:05 | N.A. | 1444 | Yes | BZD, Cisordinol, SSRI | Suicide, hanging |
| MD6 | MDD | 59 | F | 20:35 | 00:00 | 6.53 | 1199 | Yes | BZD | Suicide, suffocation |
| MD7 | MDD | 56 | F | 10:30 | 14:00 | 6.42 | 1120 | Yes | BZD | Suicide, intoxication |
| MD8 | MDD | 62 | F | 11:40 | 16:15 | 6.18 | 1155 | Yes | BZD | Suicide, strangulation |
| MD9 | MDD | 58 | M | 4:30 | 9:40 | 6.80 | 1470 | Yes | SGA, SSRI, TCA | Suicide, intoxication |
| MD10 | MDD | 90 | F | 3:55 | 11:20 | 6.62 | 1220 | No | BZD, INN | Suicide, intoxication |
| MD11 | BD | 81 | M | 6:40 | 20:00 | 6.70 | 1283 | No | BZD, Li, TCA, VPA | Legal euthanasia |
| MD12 | MDD | 77 | F | 8:40 | 11:20 | 6.77 | 1120 | Yes | BZD, INN, SSRI | Legal euthanasia |
| MD13 | MDD | 50 | M | 9:40 | 13:00 | 6.80 | 1420 | No | BZD | Legal euthanasia |
| MD14 | MDD | 38 | M | 6:15 | 11:45 | 6.79 | 1525 | No | BZD | Legal euthanasia |
| MD15 | MDD | 61 | F | 8:40 | 15:35 | 7.07 | 1205 | No | BZD | Legal euthanasia |
| MD16 | MDD | 61 | M | 5:45 | 14:05 | N.A. | 1210 | No | SSRI | Legal euthanasia |
| MD17 | MDD | 67 | M | 3:30 | 11:10 | 6.72 | 1445 | No | BZD, Hal | Legal euthanasia |
| MD18 | MDD | 70 | M | 20:00 | 19:00 | N.A. | 1500 | No | BZD, DA stabilizer, SSRI | Heart attack |
| MD19 | MDD | 73 | F | 22:00 | 19:00 | N.A. | 1287 | No | BZD, TCA | Bronchopneumonia |

| | | | | | | | | | | |
|---|---|---|---|---|---|---|---|---|---|---|
| MD20 | MDD | 71 | M | 13:30 | 22:30 | N.A. | 1109 | No | BZD, Li, MAOI, PTZ | Respiratory insufficiency |
| MD21 | MDD | 61 | M | 60:20 | 4:40 | N.A. | 1424 | No | PTZ | Pneumonia |
| MD22 | MDD | 81 | M | 6:00 | 15:30 | 6.50 | 1280 | No | Hal | Renal insufficiency |
| MD23 | MDD | 68 | M | 8:55 | 5:45 | 6.82 | 1510 | Yes | BZD, SSRI | Sudden death |
| MD24 | BD | 70 | M | 6:50 | 21:00 | 6.19 | 1275 | No | BZD, LTG, VPA | Pulmonary infection and renal insufficiency |
| MD25 | BD | 73 | M | 13:45 | 19:45 | 6.24 | 1480 | No | BZD | Infection |
| MD26 | BD | 94 | F | 5:10 | 4:00 | 7.55 | 985 | No | BZD, SSRI | Respiratory and cardiac failure |
| MD27 | MDD | 58 | F | 7:20 | 23:00 | 5.61 | 1295 | No | BZD, DA stabilizer, SSRI | Renal insufficiency |
| MD28 | BD | 65 | M | 4:50 | 18:30 | N.A. | 1305 | No | BZD, Li, SGA | Colon carcinoma |
| **Median** | - | 66 | - | 08:55 | - | 6.70 | 1291 | - | - | - |
| CTR1 | - | 68 | F | 5:45 | 12:15 | 6.97 | 1135 | No | - | Legal euthanasia |
| CTR2 | - | 83 | F | 3:20 | 16:30 | 6.80 | 1173 | No | - | Legal euthanasia |
| CTR3 | - | 49 | M | 6:15 | 17:30 | 6.15 | 1364 | No | BZD | Legal euthanasia |
| CTR4 | - | 79 | M | 6:30 | 10:00 | 6.71 | 1121 | No | BZD | Legal euthanasia |
| CTR5 | - | 83 | M | 5:45 | 19:35 | 7.28 | 1195 | No | - | Legal euthanasia |
| CTR6 | - | 41 | M | 17:00 | 00:00 | N.A. | 1150 | No | - | Renal insufficiency |
| CTR7 | - | 72 | F | 17:30 | 16:30 | N.A. | N.A. | No | - | Bronchopneumonia |
| CTR8 | - | 49 | M | 21:40 | 19:20 | N.A. | 1629 | No | - | Cardiac infarction |
| CTR9 | - | 66 | M | 41:00 | 00:00 | N.A. | 1461 | No | INN | Septic shock |
| CTR10 | - | 88 | F | 5:55 | 7:00 | 6.05 | 1115 | No | BZD | Cardiac arrest |
| CTR11 | - | 39 | M | 16:30 | 00:30 | N.A. | 1400 | No | - | Cardiac infarction |
| CTR12 | - | 55 | M | 7:15 | 11:05 | N.A. | 1393 | No | BZD, INN | Intestinal ischemia |

| | | | | | | | | | | |
|---|---|---|---|---|---|---|---|---|---|---|
| CTR13 | - | 78 | F | 4:35 | 22:40 | 6.41 | 1226 | No | BZD, INN | Bronchopneumonia |
| CTR14 | - | 73 | M | 8:00 | 16:15 | 5.37 | 1553 | No | BZD, INN | Pneumonia |
| CTR15 | - | 83 | M | 5:15 | 11:15 | 6.60 | 1372 | No | BZD | Cardiac infarction |
| CTR16 | - | 67 | M | 9:00 | 15:05 | 6.48 | 1292 | No | BZD, INN | Aortic aneurysm |
| **Median** | - | 70 | - | 6:52 | - | 6.54 | 1292 | - | - | - |
| ***p*-value** | - | 0.54 | 0.95 | 0.35 | n.sig. | 0.50 | 0.60 | - | - | - |

**Abbreviations:** BD, bipolar disorder; BW, brain weight; BZD, benzodiazepine; CTD, clock time of death; CTR, control; DA, dopamine; F, female; g, gram; h, hour; Hal, haloperidol; INN, nemifitide; Li, lithium; LTG, lamotrigine; M, male; MAOI, monoamine oxidase inhibitor; MD, mood disorders; MDD, major depressive disorder; N.A., not available; PMD, postmortem delay; PTZ, pentylenetetrazole; RIMA, reversible inhibitor of monoamine oxidase-A; SGA, second-generation antipsychotics; SSRI, selective serotonin reuptake inhibitor; TCA, tricyclic antidepressants; VPA, valproate; y, year.

**Table S1e Demographic information of Suicide-PFC collection (SMRI)**

| Diagnosis Cause of death | Cases, n | Psychiatric diagnosis (MDD/BD) | Age (y) | Sex (M/F) | PMD (h) | Brain pH | BW (g) | Psychotic features (NP/P) | Fluphenazine (mg) |
|---|---|---|---|---|---|---|---|---|---|
| **MD** | 54 | 24/30 | 44 (19-64) | 28/26 | 29 (12-84) | 6.60 (5.92-6.97) | 1440 (1170-1780) | 28/26 | 850 (0-130000) |
| Suicide | 30 | 17/13 | 42 (24-63) | 17/13 | 27.5 (13-70) | 6.61 (6.07-6.90) | 1468 (1170-1780) | 15/15 | 100 (0-15000) |
| Natural causes | 24 | 7/17 | 44.5 (19-64) | 11/13 | 30.5 (12-84) | 6.55 (5.92-6.97) | 1355 (1170-1590) | 13/11 | 2000 (130000) |
| **Ctr** | 46 | - | 45.5 (24-63) | 33/13 | 28.5 (9-58) | 6.67 (6.00-7.03) | 1413 (1120-1900) | - | - |

**Abbreviations:** NP, individuals without a history of psychotic features; P, individuals with a history of psychotic features.

**Table S1f Clinico-pathological information of patients with suicide, MD, and control subjects (Suicide-PFC collection, SMRI)**

| Sample number | Sex | Age (y) | PMD (h) | Brain pH | BW (g) | RIN | Brain Region | Psychotic feature | Fluphenazine (mg) | Cause of Death |
|---|---|---|---|---|---|---|---|---|---|---|
| MDD1 | F | 48 | 24 | 6.36 | 1330 | 8.0 | L DLPFC, L ACC | Yes | 6500 | SUIC: OD |
| MDD2 | F | 40 | 49 | 6.72 | 1450 | 8.8 | L DLPFC, L ACC | Yes | 1000 | SUIC: HANGING |
| MDD3 | F | 56 | 15 | 6.59 | 1370 | 8.4 | L DLPFC, L ACC | No | 0 | BURNS |
| MDD4 | M | 28 | 26 | 6.7 | 1780 | 6.9 | R DLPFC, R ACC | Yes | 3000 | SUIC: HANGING |
| MDD5 | M | 35 | 19 | 6.6 | 1335 | 8.2 | R DLPFC, R ACC | Yes | 2000 | SUIC: HANGING |
| MDD6 | F | 32 | 19 | 6.7 | 1280 | 8.9 | L DLPFC, L ACC | Yes | 100 | SUIC: HANGING |
| MDD7 | F | 32 | 19 | 6.8 | 1470 | 8.0 | R DLPFC, R ACC | No | 0 | SUIC: HANGING |
| MDD8 | M | 63 | 31 | 6.6 | 1540 | 7.8 | L DLPFC, L ACC | Yes | 4000 | SUIC: HANGING |
| MDD9 | F | 51 | 36 | 6.3 | 1440 | 7.3 | L DLPFC, L ACC | Yes | 700 | UNKNOWN |
| MDD10 | M | 35 | 36 | 6.6 | 1710 | 6.3 | L DLPFC, L ACC | Yes | 0 | SUIC: GSW |
| MDD11 | M | 44 | 24 | 6.52 | 1550 | 7.9 | R DLPFC, R ACC | No | 3000 | CARDIAC |
| MDD12 | M | 56 | 38 | 6.59 | 1365 | 7.3 | L DLPFC, L ACC | No | 1000 | SUIC: OD |
| MDD13 | M | 33 | 25 | 6.86 | 1640 | 7.6 | R DLPFC, R ACC | No | 0 | SUIC: HANGING |
| MDD14 | M | 34 | 24 | 6.79 | 1425 | 8.2 | R DLPFC, R ACC | No | 0 | SUIC: JUMPED |
| MDD15 | F | 45 | 29 | 6.9 | 1350 | 7.4 | L DLPFC, L ACC | No | 0 | CARDIAC |
| MDD16 | M | 53 | 21 | 6.64 | 1520 | 7.5 | L DLPFC, L ACC | No | 0 | CARDIAC |
| MDD17 | M | 62 | 65 | 6.57 | 1490 | 7.5 | R DLPFC, R ACC | No | 0 | SUIC: STABBED |
| MDD18 | F | 36 | 32 | 6.74 | 1270 | 7.3 | R DLPFC, R ACC | Yes | 2500 | PULM EMBOL |
| MDD19 | M | 40 | 52 | 6.48 | 1590 | 6.6 | L DLPFC, L ACC | Yes | 3000 | OD |
| MDD20 | F | 28 | 40 | 6.68 | 1430 | 7.1 | L DLPFC, L ACC | Yes | 0 | SUIC: OD |
| MDD21 | M | 45 | 29 | 6.75 | 1514 | 7.9 | L DLPFC, L ACC | No | 0 | SUIC: HANGING |
| MDD22 | M | 24 | 21 | 6.61 | 1737 | 7.4 | R DLPFC, R ACC | No | 0 | SUIC: OD |
| MDD23 | F | 45 | 13 | 6.58 | 1170 | 7.2 | L DLPFC, L ACC | No | 100 | SUIC: OD |
| MDD24 | F | 47 | 25 | 6.88 | 1495 | 7.7 | L DLPFC, L ACC | No | 0 | SUIC: GSW |
| BD1 | M | 29 | 48 | 6.39 | 1570 | 9.2 | R DLPFC, R ACC | Yes | 9000 | SUIC:JUMPED |
| BD2 | M | 29 | 60 | 6.7 | 1430 | 8.5 | R DLPFC, R ACC | No | 0 | SUIC:CO |

| | | | | | | | | | | |
|---|---|---|---|---|---|---|---|---|---|---|
| BD3 | M | 45 | 28 | 6.35 | 1480 | 9.1 | L DLPFC, L ACC | Yes | 10000 | CARDIAC |
| BD4 | M | 41 | 70 | 6.71 | 1625 | 7.5 | R DLPFC, R ACC | No | 0 | SUIC:OD |
| BD5 | F | 29 | 62 | 6.74 | 1330 | 7.8 | R DLPFC, R ACC | Yes | 0 | OD |
| BD6 | M | 44 | 19 | 6.74 | 1660 | 9.3 | L DLPFC, L ACC | No | 0 | SUIC:HANGING |
| BD7 | F | 48 | 18 | 6.5 | 1205 | 8.3 | R DLPFC, R ACC | No | 0 | CARDIAC |
| BD8 | M | 42 | 32 | 6.65 | 1470 | 8.4 | L DLPFC, L ACC | No | 0 | DROWNING |
| BD9 | M | 35 | 35 | 6.3 | 1490 | 7.8 | R DLPFC, R ACC | Yes | 30000 | CARDIAC |
| BD10 | F | 59 | 53 | 6.2 | 1410 | 8.5 | L DLPFC, L ACC | No | 0 | SUIC:OD |
| BD11 | M | 54 | 44 | 6.5 | 1510 | 9.2 | R DLPFC, R ACC | No | 0 | SUIC:OD |
| BD12 | F | 35 | 17 | 6.1 | 1250 | 7.9 | L DLPFC, L ACC | Yes | 3000 | SUIC:CO |
| BD13 | F | 42 | 49 | 6.65 | 1335 | 8.0 | R DLPFC, R ACC | Yes | 15000 | OD |
| BD14 | F | 58 | 35 | 6.5 | 1440 | 9.5 | R DLPFC, R ACC | Yes | 12000 | SUIC:GSW |
| BD15 | M | 64 | 16 | 6.1 | 1340 | 7.8 | L DLPFC, L ACC | Yes | 130000 | PNEUMONIA |
| BD16 | M | 59 | 84 | 6.65 | 1300 | 7.3 | L DLPFC, L ACC | No | 500 | SLEEP APNEA |
| BD17 | M | 51 | 23 | 6.67 | 1590 | 9.5 | L DLPFC, L ACC | Yes | 1200 | CARDIAC |
| BD18 | F | 63 | 32 | 6.97 | 1290 | 8.9 | R DLPFC, R ACC | No | 0 | CARDIAC |
| BD19 | F | 44 | 37 | 6.37 | 1200 | 7.3 | L DLPFC, L ACC | Yes | 30000 | MYOCARDITIS |
| BD20 | F | 56 | 26 | 6.58 | 1170 | 8.2 | R DLPFC, R ACC | No | 25000 | DROWNING |
| BD21 | F | 43 | 39 | 6.74 | 1505 | 9.3 | R DLPFC, R ACC | Yes | 4500 | SUIC:OD |
| BD22 | M | 35 | 22 | 6.58 | 1390 | 8.5 | L DLPFC, L ACC | Yes | 2000 | DROWNING |
| BD23 | F | 50 | 62 | 6.51 | 1400 | 8.3 | R DLPFC, R ACC | Yes | 15000 | SUIC:OD |
| BD24 | F | 49 | 38 | 6.39 | 1190 | 8.5 | L DLPFC, L ACC | Yes | 0 | OD |
| BD25 | F | 33 | 24 | 6.51 | 1450 | 8.6 | R DLPFC, R ACC | No | 3000 | SUIC:HANGING |
| BD26 | F | 41 | 28 | 6.44 | 1360 | 8.7 | R DLPFC, R ACC | No | 3000 | CARDIAC |
| BD27 | F | 43 | 57 | 5.92 | 1340 | 6.3 | R DLPFC, R ACC | Yes | 10000 | OD |
| BD28 | M | 56 | 23 | 6.07 | 1670 | 9.3 | L DLPFC, L ACC | Yes | 10000 | SUIC:OD |
| BD29 | M | 48 | 23 | 6.9 | 1466 | 8.4 | R DLPFC, R ACC | No | 0 | SUIC:HANGING |
| BD30 | M | 19 | 12 | 5.97 | 1484 | 7.9 | L DLPFC, L ACC | No | 2000 | OD |
| **Median** | - | 44 | 29 | 6.60 | 1440 | 8.0 | - | - | 850 | - |
| Ctr1 | M | 48 | 12 | 6.51 | 1410 | 8.6 | R DLPFC, R ACC | No | 0 | CARDIAC |

| | | | | | | | | | | |
|---|---|---|---|---|---|---|---|---|---|---|
| Ctr2 | F | 50 | 35 | 6.31 | 1520 | 7.9 | L DLPFC, L ACC | No | 0 | CARDIAC |
| Ctr3 | M | 50 | 11 | 6.5 | 1530 | 8.6 | L DLPFC, L ACC | No | 0 | CARDIAC |
| Ctr4 | M | 63 | 37 | 6.5 | 1530 | 8 | L DLPFC, L ACC | No | 0 | CARDIAC |
| Ctr5 | M | 24 | 17 | 6.6 | 1595 | 8.1 | L DLPFC, L ACC | No | 0 | MVA |
| Ctr6 | M | 44 | 27 | 6.82 | 1410 | 7.2 | R DLPFC, R ACC | No | 0 | ACUTE ALCOHOL POISONING |
| Ctr7 | M | 35 | 31 | 6.59 | 1520 | 8 | R DLPFC, R ACC | No | 0 | MVA |
| Ctr8 | M | 63 | 40 | 6.91 | 1410 | 8.2 | R DLPFC, R ACC | No | 0 | CARDIAC |
| Ctr9 | M | 34 | 9 | 6.56 | 1535 | 5.9 | R DLPFC, R ACC | No | 0 | MVA |
| Ctr10 | F | 56 | 29 | 6.78 | 1278 | 8.9 | L DLPFC, L ACC | No | 0 | CARDIAC |
| Ctr11 | F | 56 | 31 | 6.66 | 1400 | 6.1 | L DLPFC, L ACC | No | 0 | OBESITY |
| Ctr12 | F | 39 | 24 | 6.88 | 1200 | 7.3 | R DLPFC, R ACC | No | 0 | CARDIAC |
| Ctr13 | F | 44 | 28 | 6.59 | 1330 | 8.3 | R DLPFC, R ACC | No | 0 | CARDIAC |
| Ctr14 | M | 49 | 46 | 6.5 | 1605 | 8.3 | R DLPFC, R ACC | No | 0 | CARDIAC |
| Ctr15 | M | 53 | 9 | 6.4 | 1500 | 9.1 | L DLPFC, L ACC | No | 0 | CARDIAC |
| Ctr16 | M | 37 | 13 | 6.5 | 1600 | 8.3 | L DLPFC, L ACC | No | 0 | CARDIAC |
| Ctr17 | M | 51 | 31 | 6.7 | 1400 | 7.3 | R DLPFC, R ACC | No | 0 | CARDIAC |
| Ctr18 | M | 53 | 28 | 6 | 1340 | 8.4 | L DLPFC, L ACC | No | 0 | CARDIAC |
| Ctr19 | F | 38 | 33 | 6 | 1120 | 9.7 | R DLPFC, R ACC | No | 0 | CARDIAC |
| Ctr20 | F | 38 | 28 | 6.7 | 1350 | 8.8 | R DLPFC, R ACC | No | 0 | CARDIAC |
| Ctr21 | M | 60 | 47 | 6.8 | 1460 | 8.4 | R DLPFC, R ACC | No | 0 | CARDIAC |
| Ctr22 | M | 35 | 52 | 6.7 | 1700 | 8.7 | R DLPFC, R ACC | No | 0 | MYOCARDITIS |
| Ctr23 | M | 34 | 22 | 6.48 | 1480 | 8.2 | R DLPFC, R ACC | No | 0 | CARDIAC |
| Ctr24 | M | 47 | 21 | 6.81 | 1550 | 8.7 | L DLPFC, L ACC | No | 0 | CARDIAC |
| Ctr25 | M | 45 | 29 | 6.94 | 1405 | 8.5 | R DLPFC, R ACC | No | 0 | CARDIAC |
| Ctr26 | F | 34 | 24 | 6.87 | 1255 | 6.6 | R DLPFC, R ACC | No | 0 | CARDIAC |
| Ctr27 | M | 42 | 37 | 6.91 | 1340 | 7.8 | L DLPFC, L ACC | No | 0 | CARDIAC |
| Ctr28 | F | 44 | 10 | 6.2 | 1305 | 8.4 | R DLPFC, R ACC | No | 0 | CARDIAC |
| Ctr29 | M | 45 | 18 | 6.81 | 1585 | 8.0 | L DLPFC, L ACC | No | 0 | CARDIAC |
| Ctr30 | M | 49 | 23 | 6.93 | 1390 | 9.5 | L DLPFC, L ACC | No | 0 | CARDIAC |

| | | | | | | | | | | |
|---|---|---|---|---|---|---|---|---|---|---|
| Ctr31 | M | 32 | 24 | 7.03 | 1415 | 8.3 | L DLPFC, L ACC | No | 0 | CARDIAC |
| Ctr32 | M | 55 | 31 | 6.7 | 1515 | 8.3 | L DLPFC, L ACC | No | 0 | CARDIAC |
| Ctr33 | F | 49 | 45 | 6.72 | 1435 | 8.6 | L DLPFC, L ACC | No | 0 | CARDIAC |
| Ctr34 | F | 33 | 29 | 6.52 | 1360 | 8.3 | L DLPFC, L ACC | No | 0 | ASTHMA |
| Ctr35 | M | 48 | 31 | 6.86 | 1580 | 7.0 | R DLPFC, R ACC | No | 0 | CARDIAC |
| Ctr36 | M | 50 | 49 | 6.75 | 1645 | 8.0 | R DLPFC, R ACC | No | 0 | CARDIAC |
| Ctr37 | M | 32 | 13 | 6.57 | 1410 | 9.0 | R DLPFC, R ACC | No | 0 | CARDIAC |
| Ctr38 | M | 47 | 11 | 6.6 | 1495 | 8.3 | L DLPFC, L ACC | No | 0 | CARDIAC |
| Ctr39 | M | 46 | 31 | 6.67 | 1360 | 8.6 | L DLPFC, L ACC | No | 0 | CARDIAC |
| Ctr40 | M | 40 | 38 | 6.67 | 1498 | 8.7 | L DLPFC, L ACC | No | 0 | CARDIAC |
| Ctr41 | M | 51 | 22 | 6.71 | 1900 | 8.7 | L DLPFC, L ACC | No | 0 | CARDIAC |
| Ctr42 | M | 31 | 11 | 6.13 | 1335 | 8.4 | R DLPFC, R ACC | No | 0 | PULM EMBOL |
| Ctr43 | M | 48 | 24 | 6.91 | 1321 | 8.6 | R DLPFC, R ACC | No | 0 | CARDIAC |
| Ctr44 | F | 39 | 58 | 6.46 | 1260 | 8.4 | R DLPFC, R ACC | No | 0 | CARDIAC |
| Ctr45 | M | 47 | 36 | 6.57 | 1535 | 8.0 | L DLPFC, L ACC | No | 0 | CARDIAC |
| Ctr46 | F | 41 | 50 | 6.17 | 1290 | 7.3 | R DLPFC, R ACC | No | 0 | CARDIAC |
| **Median** | - | 45.5 | 28.5 | 6.67 | 1413 | 8.3 | - | - | - | - |
| ***p*-value** | 0.04 | 0.62 | 0.18 | 0.16 | 0.80 | 0.21 | 0.69 | - | - | - |

**Abbreviations:** ACC, anterior cingulate cortex; BD, bipolar disorder; BW, brain weight; CO, carbon monoxide intoxication; Ctr, control; DLPFC, dorsolateral prefrontal cortex; F, female; GSW, gunshot wound; L, left; M, male; MVA, motor vehicle accident; OD, overdose drugs; PMD, postmortem delay; PULM EMBOL, pulmonary embolism; R, right; RIN, RNA integrity number; SUIC, suicide.

**Sample preparation**

**Tissues for immunohistochemical and immunofluorescent staining** Tissues from the hippocampi and hypothalami were dissected at autopsy, fixed, dehydrated, cleared, and embedded in paraffin. Coronal serial sections (6 μm) were made from the anterior or middle hippocampus and hypothalamus from the level of the lamina terminalis to mamillary bodies. Every 100$^{th}$ section from the hypothalamus and one section from the hippocampus were collected and stained with thionine for orientation. One section from the hippocampus was collected and stained with hematoxylin for cell nucleus counting in the dentate gyrus.

**Tissues for real-time PCR:** Prepared by the SMRI.

**Immunohistochemical staining** Antibodies used in this study, their manufacturers, specificities, and staining protocols are presented in **Table S2**. In brief, after deparaffinization and rehydration, the sections were washed, antigen retrieved by microwave, and/or pre-incubated to block background staining. Subsequently, the sections were incubated with primary antibody at 4 °C, then with biotinylated secondary antibody (1:400; Vector Labs, Inc. Burlingame, USA), followed by Avidin-Biotin-peroxidase Complex (1:800; Vector Labs, Inc. Burlingame, USA). Finally, sections for bright field imaging were incubated in a solution of 0.5 mg/mL 3,3-diaminobenzidine (Sigma) in a total volume of 15 mL TBS containing 5 μL $H_2O_2$ 30% (Merck) and 0.035 g ammonium Nickel sulfate (DAB-Ni), at room temperature for color development. For double labeling (DAB-Ni vs DAB), ammonium nickel sulfate was not included in the second DAB development. The enzyme reaction was stopped in aqua dest. Subsequently, the sections were dehydrated, cleared, and coverslipped in Entellan (Merck). After the incubation of primary antibodies, sections for confocal imaging were incubated with fluorescent-conjugated antibodies to visualize the immunoreactions. Finally, sections were incubated with Hoechst to visualize the nuclei and coverslipped with antifade mounting medium (VECTASHIELD) and sealed.

**Table S2a Specification of the antibodies used**

| Antibody | Species | Manufacturer | Catalog number |
|---|---|---|---|
| P2RX7 | Goat polyclonal | Novus Biologicals | NBP1-37775 |
| P2RY12 | Rabbit polyclonal | Sigma-Aldrich | HPA014518 |
| GFAP | Rabbit polyclonal | DAKO | GA524 |
| CD68 | Mouse monoclonal | DAKO | M0814 |

**Note:** 1 Specificity of commercial antibodies was provided by the manufacturers.
**Abbreviations:** CD68, the cluster of differentiation 68; GFAP, glial fibrillary acidic protein; P2RX7, P2X purinoceptor 7; P2RY12; P2Y purinoceptor 12.

**Table S2b Immunohistochemistry for single staining**

| Antibody | Antigen retrieval | Blocking buffer | Dilution |
|---|---|---|---|
| P2RX7 | 0.01M citrate buffer (pH 6.0) microwave 800 w 10 min | - | 1:800 |
| P2RY12 | 0.01M citrate buffer (pH 6.0) microwave 800 w 10 min | - | 1:2000 |
| GFAP | - | 1×TBS-4% milk | 1:100000 |
| CD68 | 0.05M Tris-HCl buffer (pH 9.0) microwave 800 w 20 min | - | 1:400 |

**Note:** SUMI: supermix, 0.25 g gelatin in 100 ml 1×TBS, pH 7.6 heat the gelatin, until it dissolves, and add Triton-X 0.5 ml. Store in fridge.

**Table S2c Immunohistochemistry for confocal staining**

| Antibody | Antigen retrieval | Primary antibodies dilutions | Secondary antibodies | Nuclear staining |
|---|---|---|---|---|
| P2RY12 | 0.05M Tris-HCl buffer (pH 9.0) microwave 800 w 20 min | 1:500 | Alexa 594-conjugated goat anti-rabbit (1:200) | Hoechst 1:1200 in SUMI 10 min |
| CD68 | | 1:100 | Alexa 488-conjugated goat anti-mouse (1:200) | |

### Image analysis

**Cell counting.** To determine the total nuclei/cells in the target areas, a tissue scanner (ZEISS Axio Scan.Z1 (Oberkochen, Germany)) was used to collect images with hematoxylin/ immunohistochemical staining at a Plan_Apochromat 20×/0.8 objective. Analysis was done with the open software QuPath 0.3.0 [2]. For the nuclei and astrocytes counting, 20% of the manually outlined areas of interest (AOI) were systematic random sampled. Following P2RX7 immunohistochemical staining, target cells (nuclei or astrocyte cell bodies with positive staining) within the AOI were quantified. Subjects with over 5% of total nuclei with P2RX7 nuclear staining were recorded, in which P2RX7$^+$ nuclei were at detectable levels in microscopy practice. In the DG, the ratio (%) of P2RX7$^+$ nuclei to total nuclei in one subject was calculated, because the granule cells were tightly packed together in a laminated and homogeneous manner. To determine the density of P2RX7$^+$ nuclei in the DG, CA1, CA2-4, Sub, and Ent, we divided the P2RX7$^+$ nuclei by the volumes of interest.

**Quantification of protein expression.** Quantitative image analyses were carried out by one investigator who was blind to the experimental conditions. The setup consisted of an image analysis system (Image-Pro version 6.3, Media Cybernetics, Rockville, USA) connected to a black and white camera (SONY XC-77E) mounted on microscopes (ZEISS Axioskop with Plan-NEOFLUAR ZEISS objectives, Carl Zeiss GmbH, Jena, Germany) or Zen system. The part of the hypothalamic nuclei and hippocampal subfields containing P2RX7, P2RY12, GFAP, or CD68 immunoreactive cells was delineated manually at a 20×/0.5 objective, according to the thionine staining. A computer program developed in-house was used to mask and extract the P2RX7, P2RY12, GFAP, or CD68 immunoreactive positive signal structures. The threshold for a positive signal was set at 0.1. The computer determined the optical density (OD) of the pixels and the percentage of surface area covered by the signal (area mask). In the hippocampal subregions, the integrated optical density (IOD) of every AOI was then calculated by multiplying the OD by the area mask corrected for background and total area. Regarding the thionine staining throughout the hypothalamic nuclei, 5 sections, including the first, first quarter, middle, last quarter, and last sections that contained PVN/SON neurons, were mounted for immunohistochemical staining and IOD quantification. The IOD of each section was calculated by multiplying the OD by the area of the mask. The IOD of each subject was calculated by adding up the IODs of five sections, then dividing by the added total AOIs of five sections. The volume of PVN/SON-containing neurons shown by thionine staining was calculated based on the Cavalieri principle [3]. Of note, in the imaging processing for SON volume quantification, AOIs from images that showed both medial and lateral SONs were delineated and added to the area of the corresponding level.

**Confocal microscopy and imaging processing.** A confocal laser-scanning microscope SP8 (DM6000 CFS; acquisition software, Leica Application Suite AF v3.2.1.9702, RRID: SCR_013673) was used for image capture. Z-stack images were collected with a 63×/1.4 oil objective at 3.5× zoom and z-step of 0.5 μm. To validate the integrity of microglial debris, which was structurally disconnected by processes, the entire P2RY12$^+$ microglia debris was captured within the z-stack for image processing. Three-dimensional reconstruction of P2RY12$^+$ microglia was performed using Imaris software (version 8.3, Bitplane AG, Zurich, Switzerland).

**Morphological parameters of P2RY12$^+$ microglia.** Using ImageJ, the total number of P2RY12$^+$ microglia in the mEnt was manually counted within the outlined area when the cell nucleoli were exposed. Results were shown in cell body number/mm$^3$. For cell size measurement, positive particles larger than 50 μm$^2$ and smaller than 400 μm$^2$ (size determined in the pilot study) were considered microglial cell body + processes. The radius of a microglia cell body was the semidiameter that could draw a max circle within the particles, resulting in pixels. For fragment measurement, positive particles smaller than 50 μm$^2$ (size determined in the pilot study) were considered microglial fragments. The size was presented as the average value per subject, resulting in average size ($\mu m^2$). The total number of microglial fragments was recorded within the outlined area. Fragment density was shown in fragment number/$mm^3$.

**Quantitative real-time PCR**

RNA isolation and cDNA synthesis were performed as described previously [4]. RNA integrity value (RIN), an indicator of human post-mortem tissue RNA quality (Stan et al., 2006), did not show any significant difference between the diagnostic groups in the SMRI material (RIN value of the ACC/DLPFC from MD: 8.06 ± 0.79 and the control group: 8.09 ± 0.74, mean ± SEM). A cDNA template (equivalent to 5 ng of total RNA) was amplified in a final volume of 10 µl using SYBR Green PCR master mix (Applied Biosystems, CA, USA) and a mixture of forward and reverse primers (each 2 pmoL/µl).

Data were acquired and processed automatically by the Applied Biosystems 7300 Real-Time PCR System. The specificity of amplification was checked by melting curve analysis. Sterile water and RNA samples without the addition of reverse transcriptase during cDNA synthesis served as negative controls. The linearity of each qPCR assay was tested by preparing a series of dilutions of the same stock cDNA in multiple plates. To reduce the effect of sample variability, reference genes were selected based on expression stability measurement [5] and the proportion of explained variability regarding all target genes. The higher stability of a reference gene means that its variance was relatively lower than the other considered reference genes. Further, if the expression of a reference gene reflects the variability of the samples well, it is expected that its application should reduce the overall variability of the target genes. Briefly, we used the following reference genes: actin beta (ACTβ), glyceraldehyde-phosphate dehydrogenase (GAPDH), hypoxanthine phosphoribosyltransferase 1 (HPRT1), tubulin alpha (TUBα), tubulin beta (TUBβ), and ubiquitin C (UBC). For primer information see **Table S3**.

**Table S3 Reference gene and target gene sequence**

| Gene | Full name | Accession Code | Forward Primer | Reverse Primer |
|---|---|---|---|---|
| **Reference Gene** | | | | |
| ACTβ | Actin beta | NM_001101 | CCCAGCCATGTACGTTGCTA | TCACCGGAGTCCATCACGAT |
| GAPDH | Glyceraldehyde-phosphate dehydrogenase | NM_002046 | CAAATTCCATGGCACCGTC | TCTCGCTCCTGGAAGATGGT |
| HPRT1 | Hypoxanthine phosphoribosyltransfe rase 1 | NM_000194 | GGACAGGACTGAACGTCTTGC | ATAGCCCCCCTTGAGCACAC |
| TUBα | Tubulin alpha | NM_006082 | CTTTGAGCCAGCCAACCAGA | GTACAACAGGCAGCAAGCCAT |
| TUBβ | Tubulin beta | NM_006087 | GGGCCAAGTTTTGGGAGGT | CACTGTCCCCATGGTATGTGC |
| UBC | Ubiquitin C | NM_021009 | GCTGCTCATAAGACTCGGCC | GTCACCCAAGTCCCGTCCTA |
| **Target Gene** | | | | |
| P2RX7 | P2X purinoceptor 7 | NM_002562 | CGTGGAGAATGGAGTGAAGA | GGGATACTCGGGACACAA |

**Figure S1**

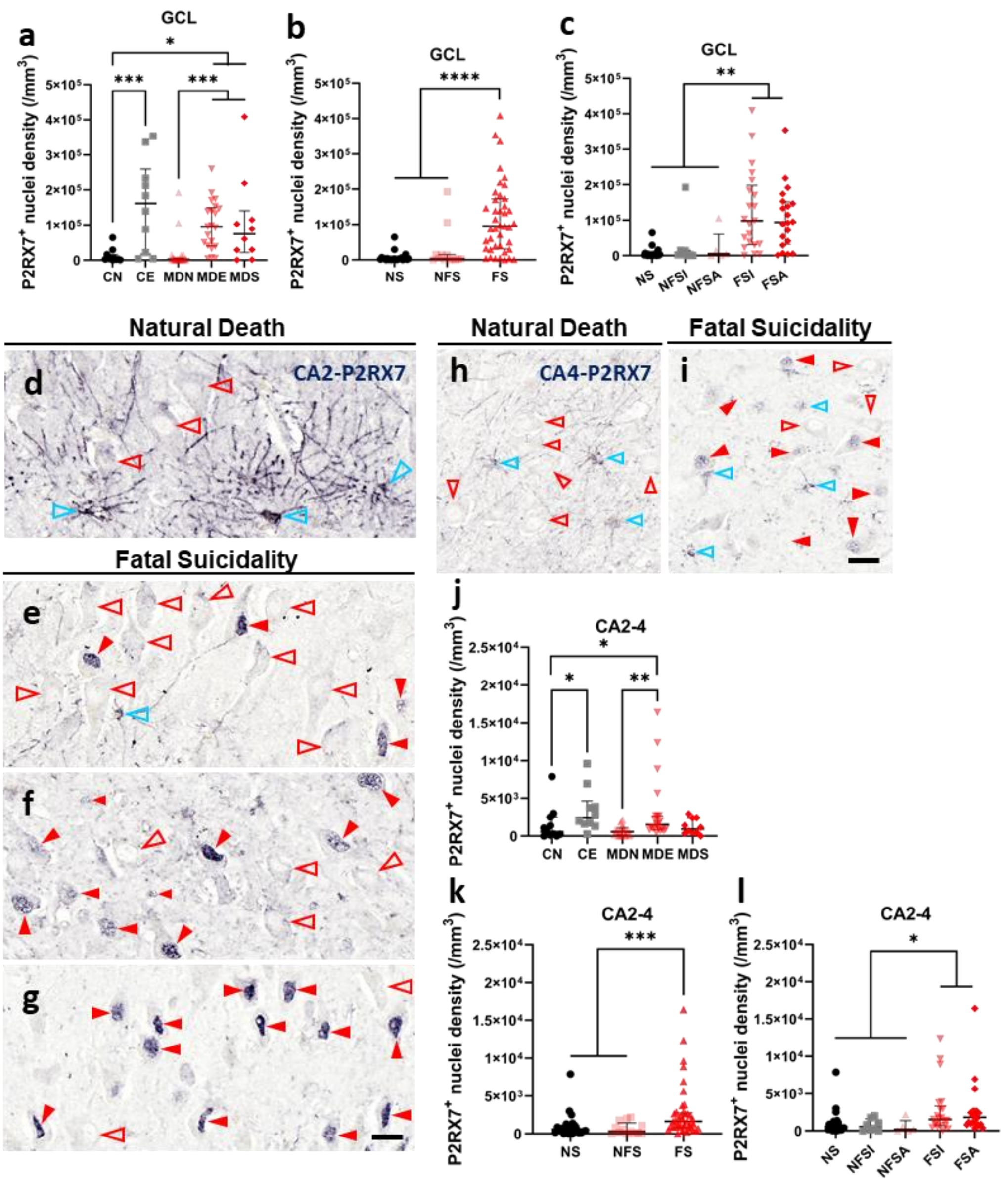


**Fig. S1 P2RX7⁺ nuclei in the DG and CA2-4 mark fatal suicidality, irrespective of MD. a-c** show the P2RX7⁺ nuclei densities in the GCL in accordance with P2RX7⁺ nuclei ratios presented in **Fig. 1i-k**. **d-l** Nuclear P2RX7 expression in the CA2 and CA4 of individuals without (**d** and **h**) and with fatal suicidality (**e-g** and **i**). Red solid arrowheads point to neurons with P2RX7⁺ nuclei. Red empty arrowheads point to neurons without nuclear P2RX7 staining. Blue empty arrowheads point to astrocytes. **j-l** More nuclei in the CA2-4 expressed P2RX7 in both Ctr and MD in relation to fatal suicidality. Scale bars, 20 μm. Data are presented as medians with interquartile ranges. * indicates $0.01 \leqslant p < 0.05$, ** indicates $0.001 \leqslant p < 0.01$, *** indicates $0.0001 \leqslant p < 0.001$.

### Figure S2

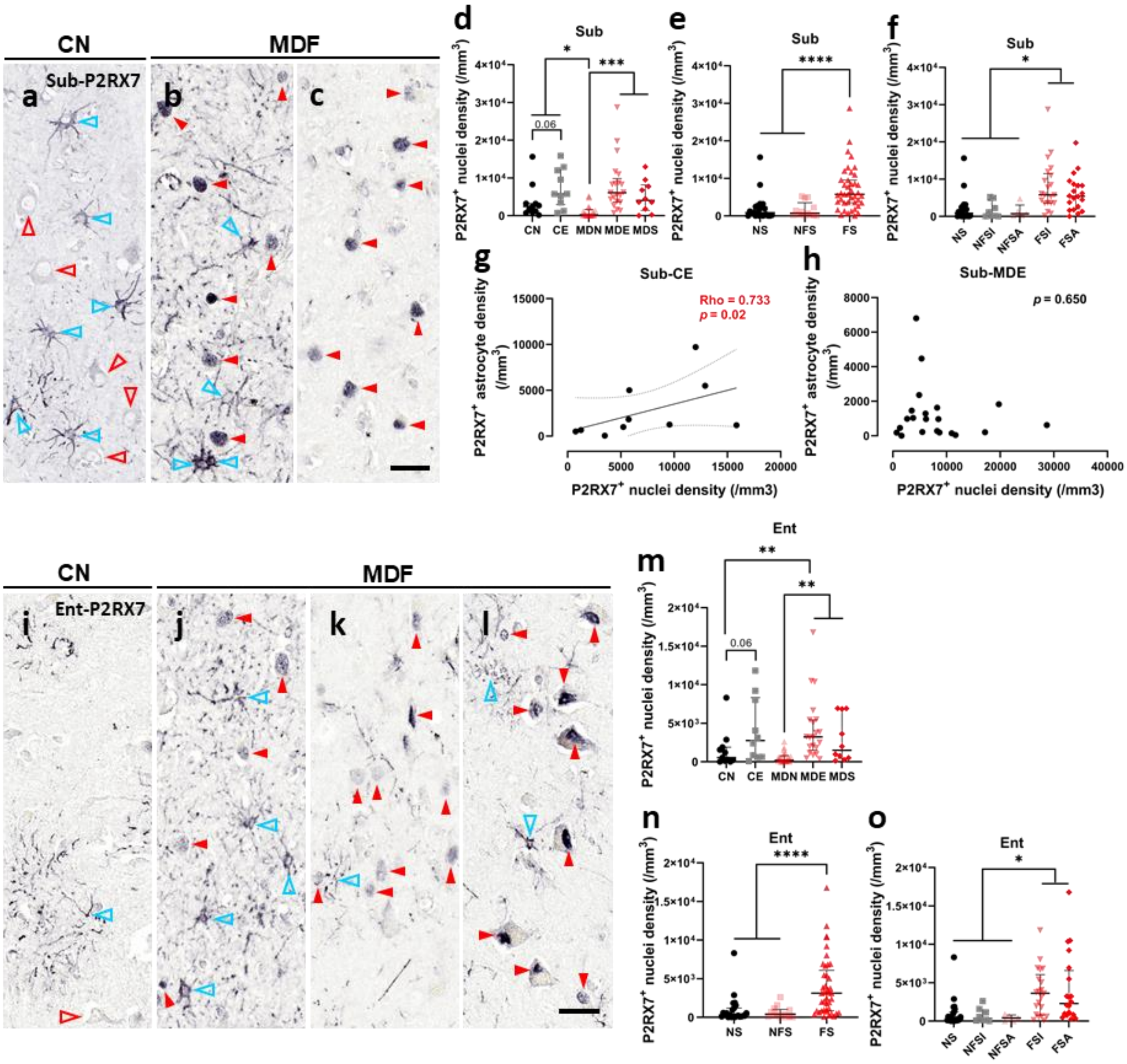


**Fig. S2 P2RX7$^+$ nuclei in the Sub and Ent mark fatal suicidality in MD.** P2RX7 expression in the Sub (**a-h**) and Ent (**i-o**) of individuals without (**a** and **i**) and with fatal suicidality (**b** and **c**, **j-l**). Red solid arrowheads point to neurons with P2RX7$^+$ nuclei. Red empty arrowheads point to neurons without nuclear P2RX7 staining. Blue empty arrowheads point to astrocytes. **d-f** More nuclei in the Sub express P2RX7 in MD in relation to fatal suicidality, which is a trend of increase within the control subsets. **g** shows a positive correlation between P2RX7$^+$ nuclei and P2RX7$^+$ astrocytes in the Sub of control subjects who died by legal euthanasia, which is absent in the MD subjects who died of the same cause (**h**). **m-o** More nuclei in the Ent express

P2RX7 in MD in relation to fatal suicidality, which is an increasing trend in CE versus CN. Scale bars, 20 μm. Data are presented as medians with interquartile ranges. * indicates $0.01 \leqslant p < 0.05$, ** indicates $0.001 \leqslant p < 0.01$, *** indicates $0.0001 \leqslant p < 0.001$.

Figure S3

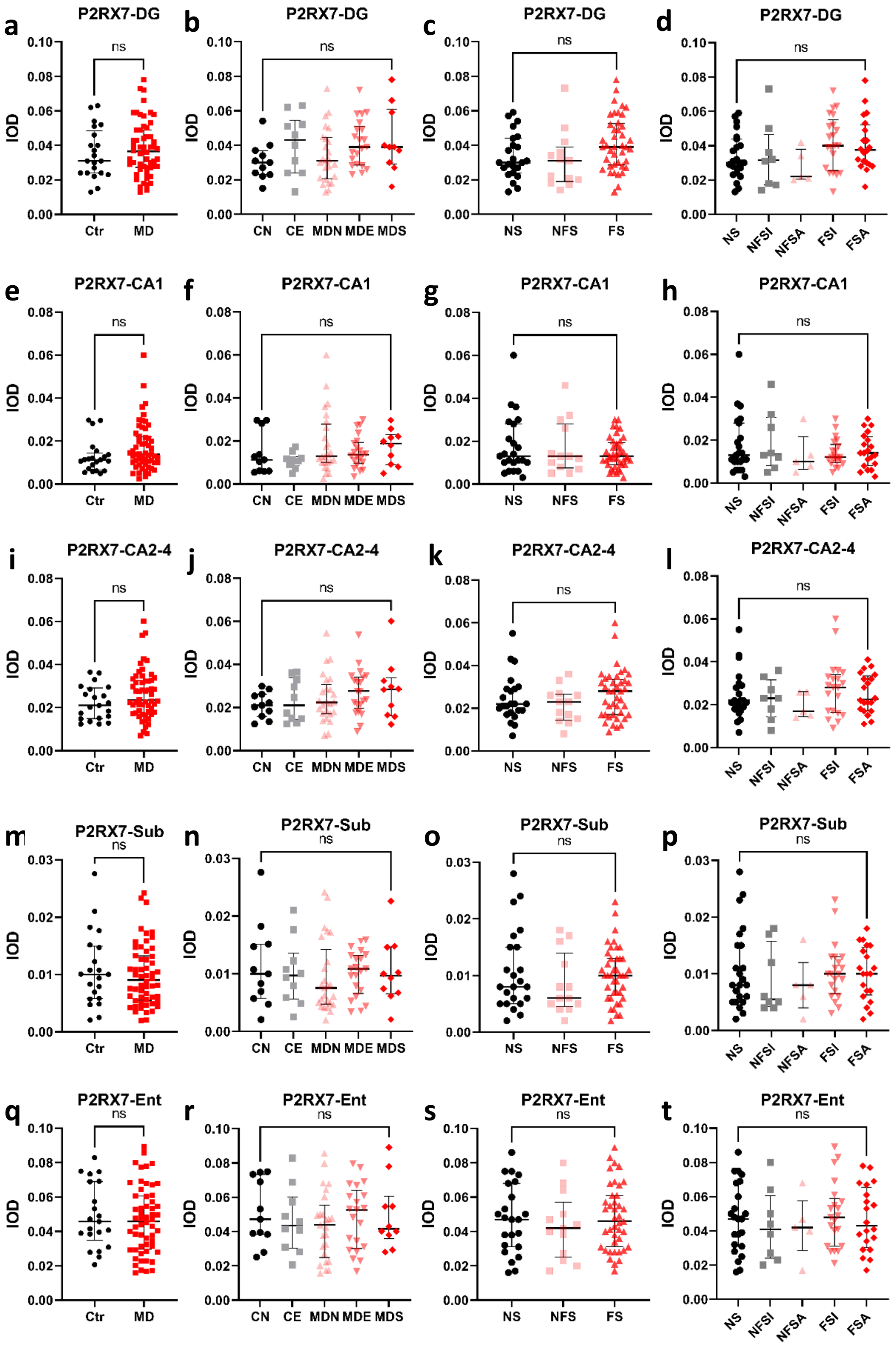


**Fig. S3 Unchanged P2RX7 total expression throughout the hippocampal subfields in relation to suicide or MD.** Data are presented as medians with interquartile ranges. ns, not significant.

Figure S4

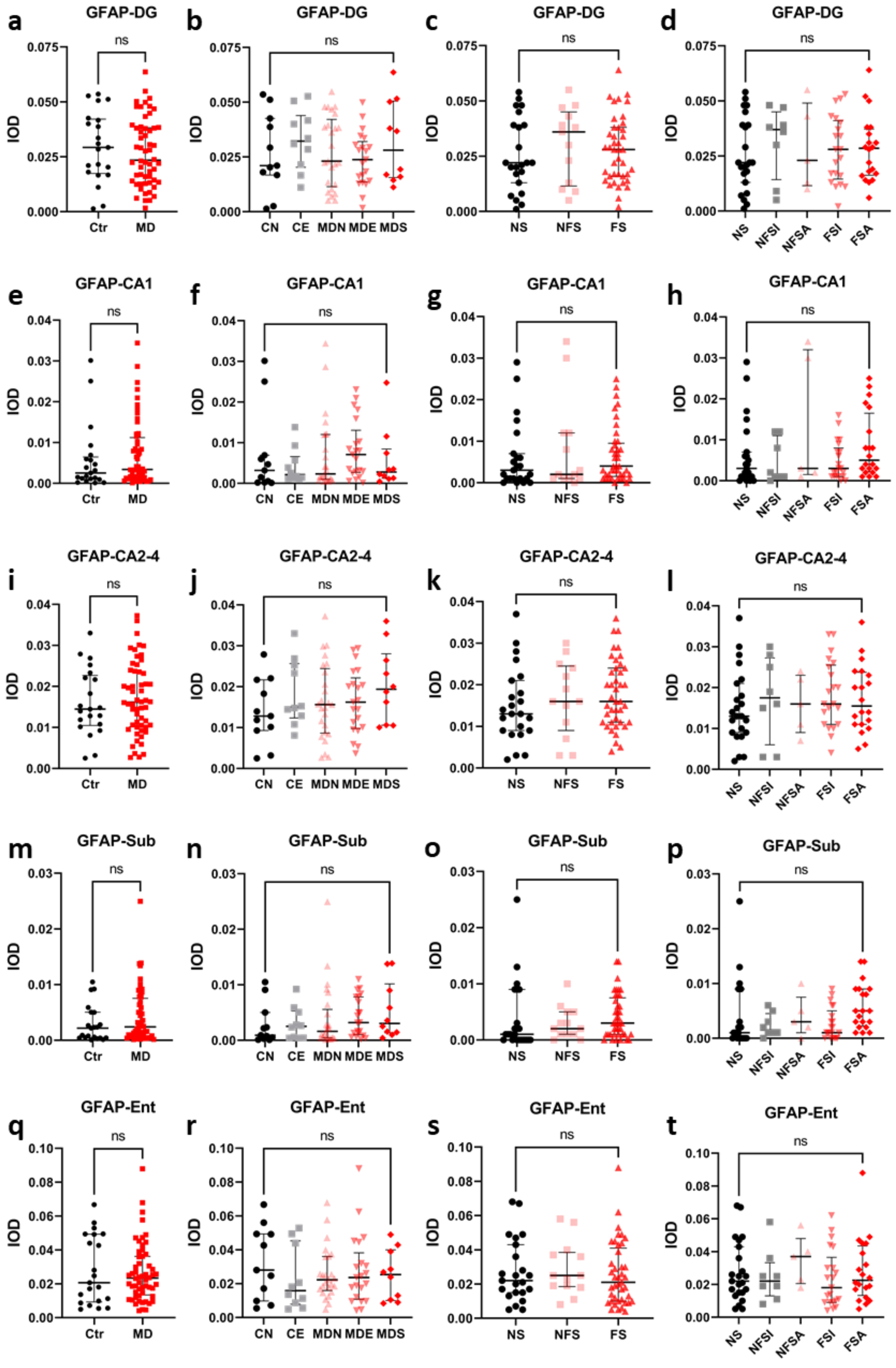


**Fig. S4 Unchanged GFAP total expression throughout the hippocampal subfields in relation to suicide or MD.** Data are presented as medians with interquartile ranges.

**Figure S5**

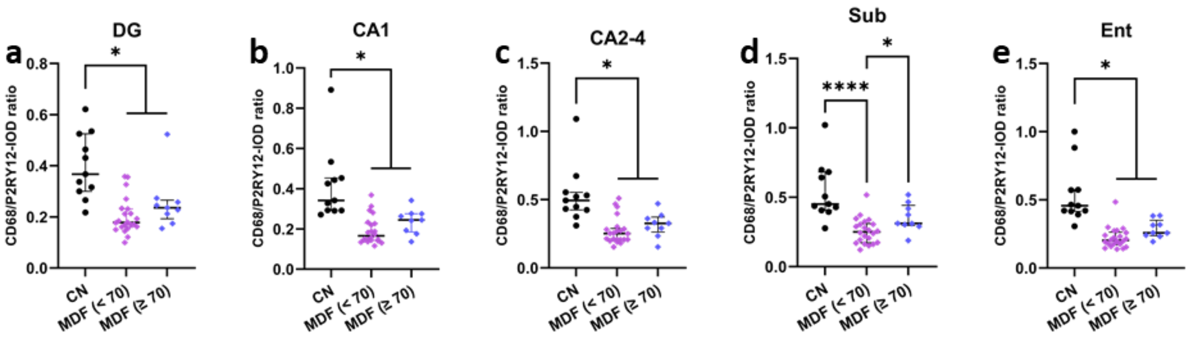


**Fig. S5 Age did not confound the P2RY12$^+$ microglial priming in the hippocampal subareas except for the Sub. a-e** The ratios of CD68 to P2RY12 from depressed patients who had fatal suicidality (MDF) are subdivided by the age of 70 and compared with non-suicidal controls. Data are presented as medians with interquartile ranges. * indicates $0.01 \leqslant p < 0.05$, **** indicates $p < 0.0001$.

### Figure S6

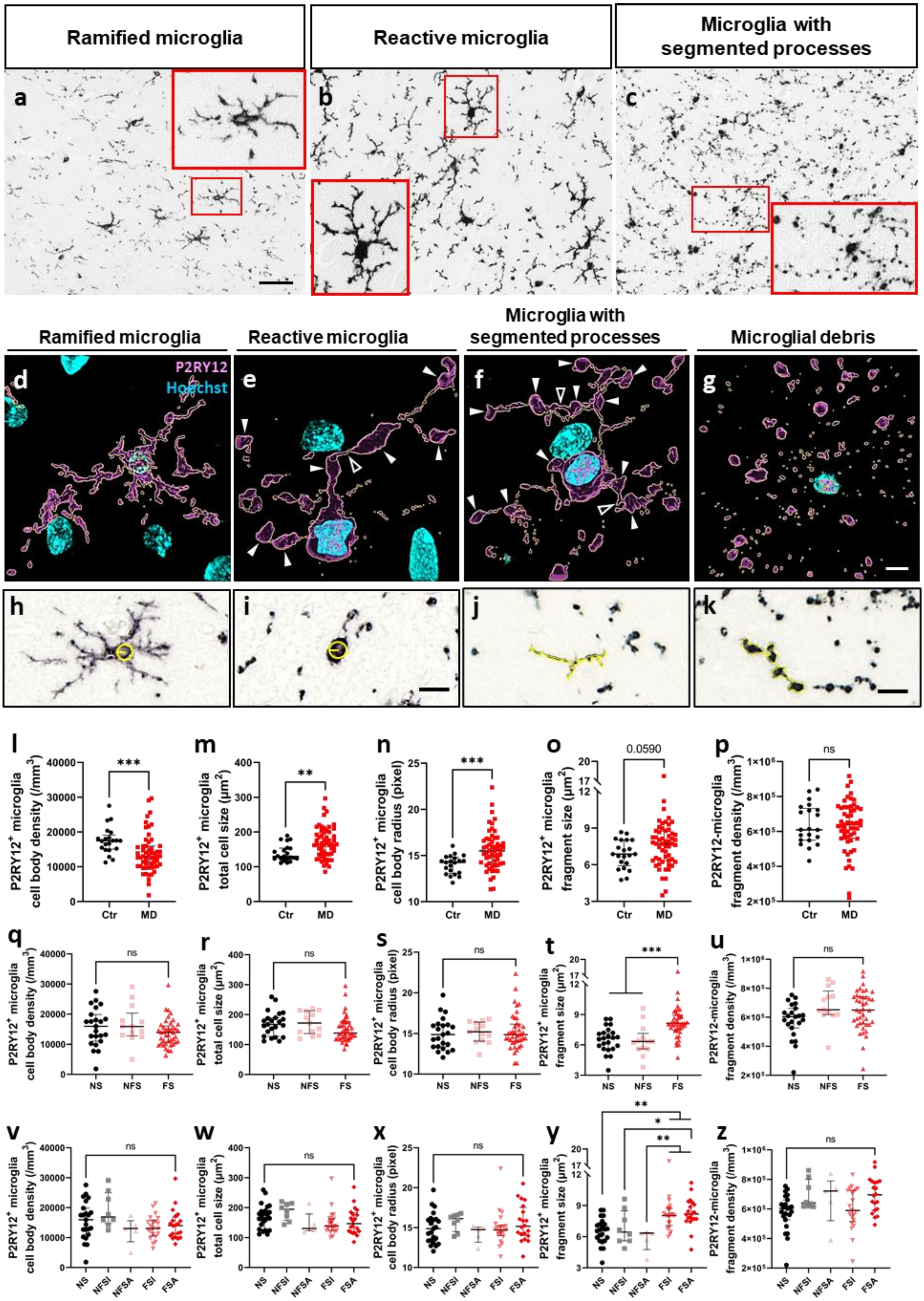

Ramified microglia
Reactive microglia
Microglia with segmented processes
a
b
c
Ramified microglia
Reactive microglia
Microglia with segmented processes
Microglial debris
d
P2RY12
Hoechst
e
f
g
h
i
j
k
l
P2RY12+ microglia cell body density (/mm3)
Ctr
MD
***
m
P2RY12+ microglia total cell size (µm2)
**
n
P2RY12+ microglia cell body radius (pixel)
***
o
P2RY12+ microglia fragment size (µm2)
0.0590
p
P2RY12-microglia fragment density (/mm3)
ns
q
r
s
t
u
NS
NFS
FS
v
w
x
y
z
NS
NFSI
NFSA
FSI
FSA

**Fig. S6 P2RY12$^+$ microglial deformation in the medial entorhinal cortex in MD and suicide.** **a-c** show ramified microglia (**a**) in a control subject who died of natural causes, reactive microglia (**b**) in a depressed patient who died of natural causes, and microglia with segmented processes (**c**) in a subject with fatal suicidality. The red boxes magnify their typical morphology. **d-g** 3D reconstruction validates process connection/disconnection of microglia phenotypes: **d** ramified microglia; **e** reactive microglia; **f** microglia with segmented processes and disconnected debris; **g** microglial microspheres and debris. White solid arrows point to the microglial microspheres. White empty arrows point to the connected processes. **h-k** graphically illustrate the measurement of microglial cell body radius and fragment size in a control case (**h** and **j**) and a suicidal case (**i** and **k**). Quantitative analyses reveal that the density of P2RY12$^+$ microglia cell bodies in subjects with MD is lower than in non-psychiatric controls (**l**). **m** and **n** P2RY12$^+$ microglia in the depressed patients have larger total cell size and cell body radius than the controls, which is a trend of increase in measuring the microglial fragments (**o**). **q-s** and **v-x** show that the sole presence of suicidality does not confound the morphological profile of the microglial cell body. **t** and **y** indicate that microglial fragments in the mEnt are larger in individuals who had fatal suicidality compared to subjects without suicidality or nonfatal suicidality. However, the density of microglial fragments in this region does not change in relation to MD (**p**), the presence of suicidality (**u**), or suicidal classifications (**z**). Scale bar, **a** 20 μm; **g** 5 μm; **i** and **k** 10 μm. Data are presented as medians with interquartile ranges. * indicates $0.01 \leqslant p < 0.05$, ** indicates $0.001 \leqslant p < 0.01$, *** indicates $0.0001 \leqslant p < 0.001$.

Figure S7

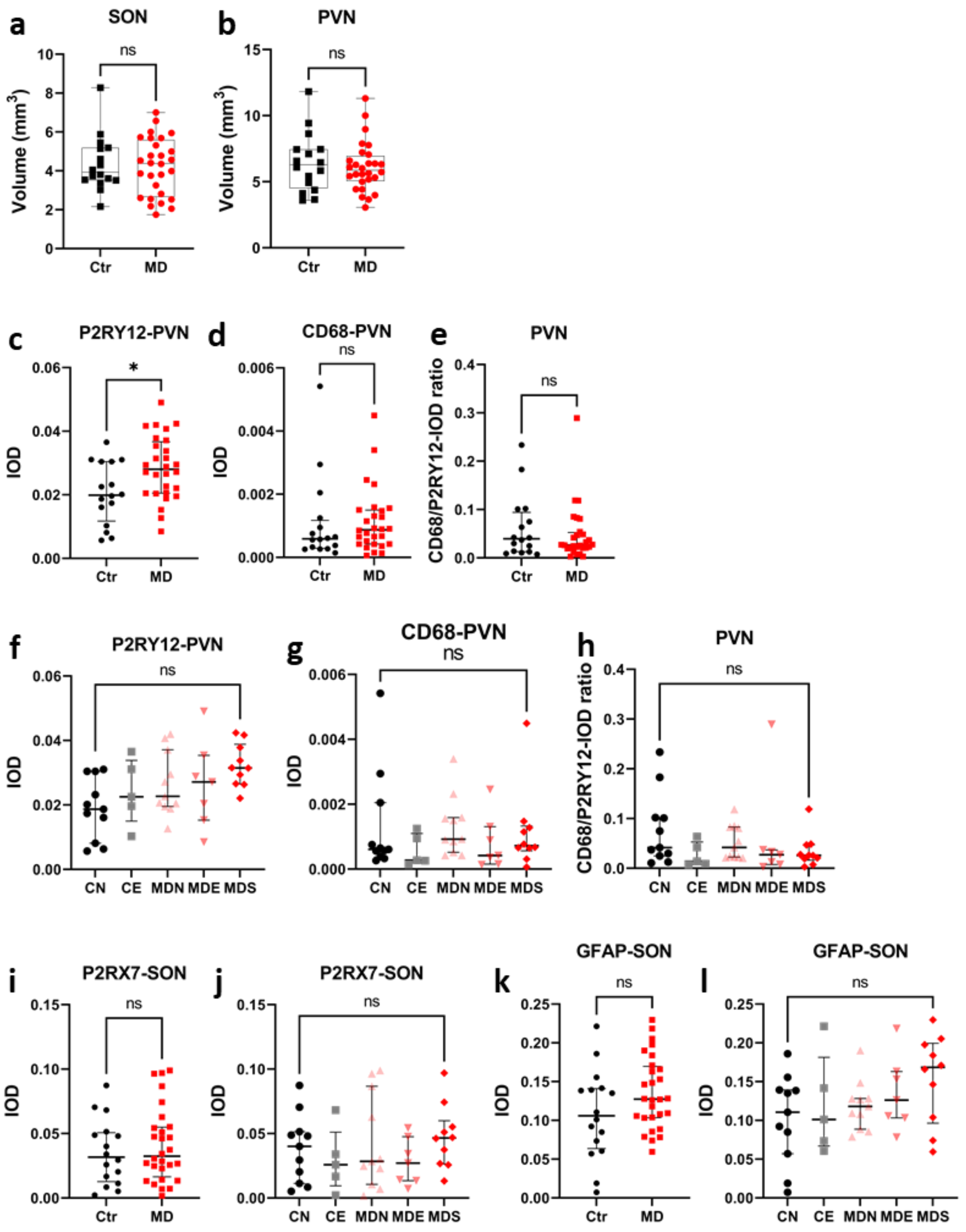


**Fig. S7 Purinergic receptors and glial expression in stress-reactive hypothalamic nuclei.** No difference in the volume of SON (**a**) or PVN (**b**) is found between Ctr and MD. **c-h** P2RY12 and CD68 expression in the PVN. **i-l** P2RX7 and GFAP expression in the SON. Data are presented as medians with interquartile ranges. * indicates $0.01 \leqslant p < 0.05$.

**Figure S8**

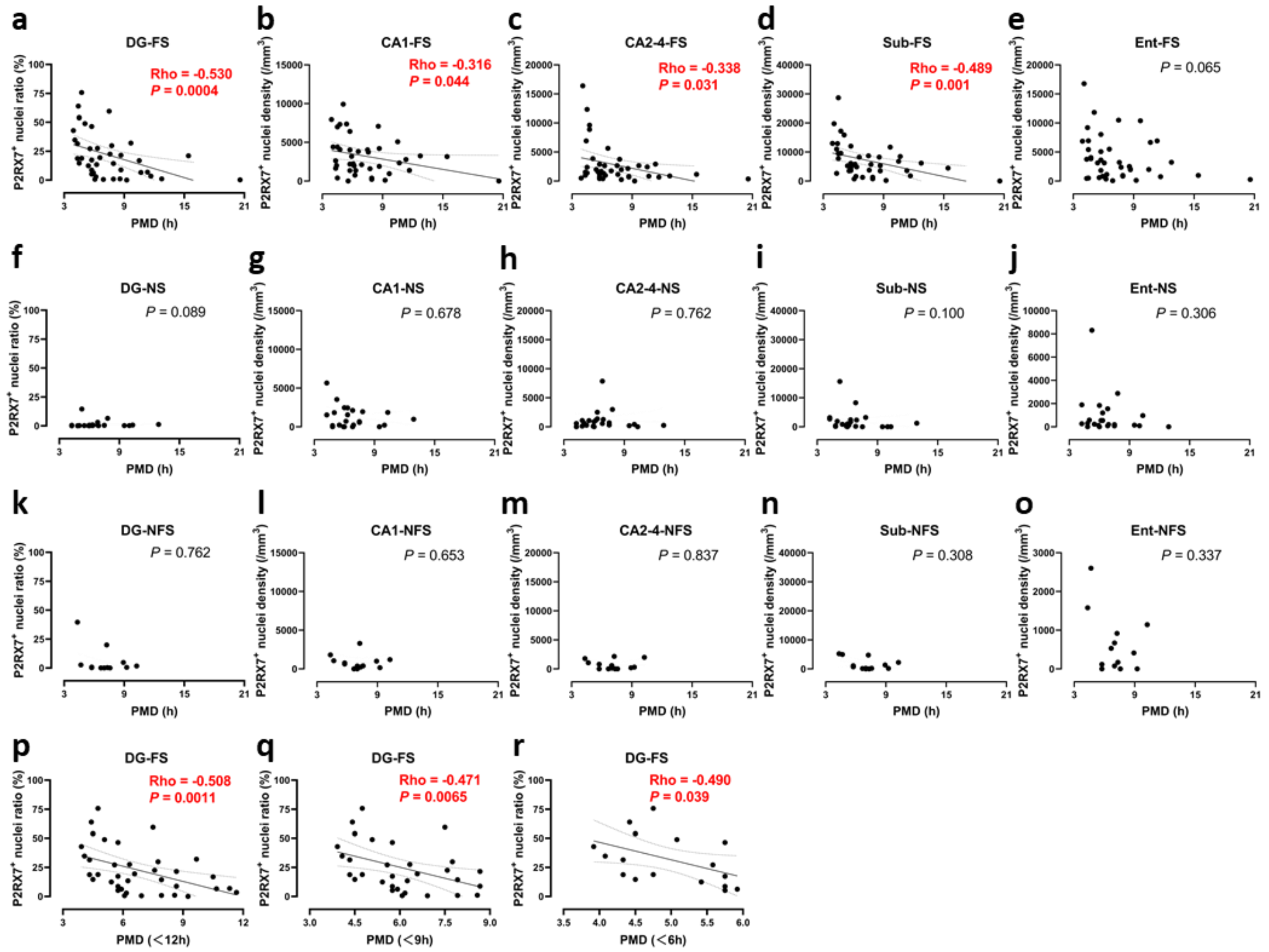


**Fig. S8 Confounder analysis of PMD.** The ratios and densities of P2RX7$^{+}$ nuclei in the hippocampal subareas (DG, CA1, CA2-4 and Sub) of individuals who had fatal suicidality is negatively associated with the PMDs (**a-e**), which is absent in individuals who did not have suicidality or nonfatal suicidality (**f-o**). **p-r** In the DG, the negative correlations remain significant when the PMDs are shortened to 12h, 9h, and 6h, respectively.

### Figure S9

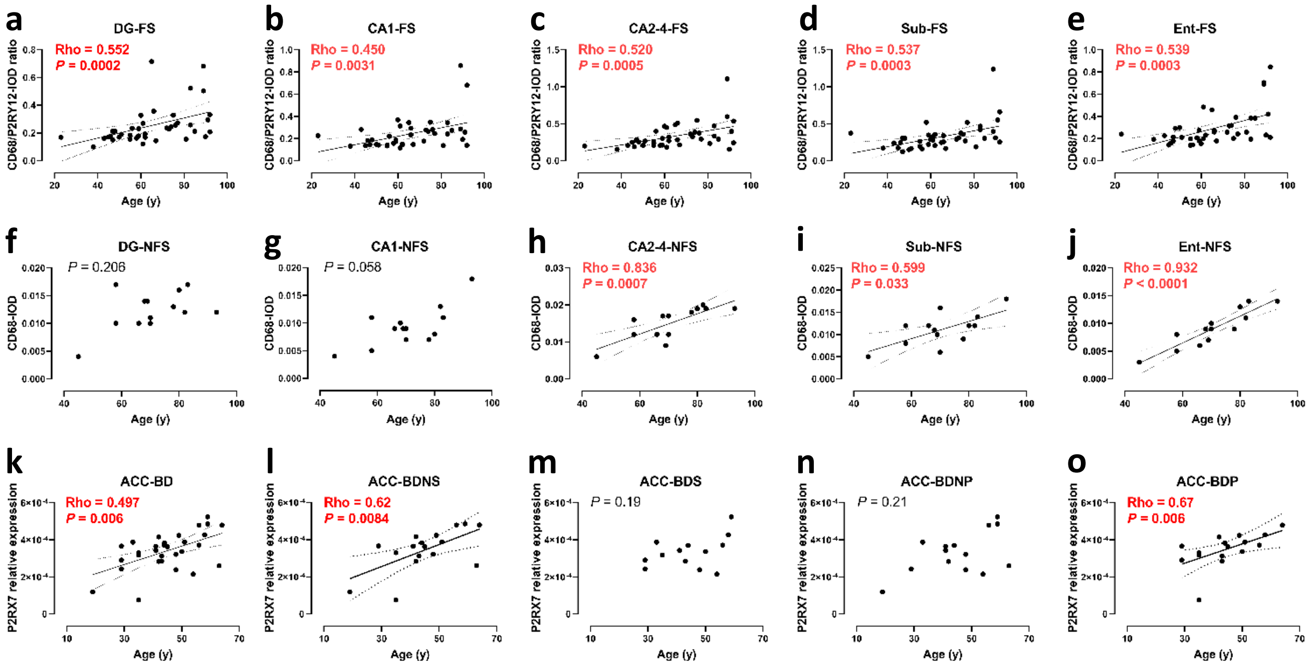


**Fig. S9 Confounder analysis of age. a-e** In individuals with fatal suicidality, the ratio of CD68 to P2RY12 throughout the hippocampal formation positively correlates with age. **f-j** In individuals with non-fatal suicidality, CD68 expression in the CA2-4, Sub and Ent shows positive correlations with age. **k-o** P2RX7 mRNA expressed through the ACC of BD positively correlates with age, which is contributed by patients with BD who developed psychotic features.
